# Functional Diversification of Gene Duplicates under the Constraint of Protein Structure

**DOI:** 10.1101/2024.03.06.583810

**Authors:** Kangning Guo, Yuqing Yang, Tingting You, Kangli Zhu

## Abstract

Gene differentiation following duplication plays a crucial role in evolution, driving the emergence of new functional genes. This process involves changes in key gene traits, such as protein structure, expression patterns, subcellular localization, and enzymatic activity, which together contribute to the development of novel functions. Here, we identified a group of homologous Glycoside Hydrolase Family 50 (GH50) agarases in the deep-sea bacterium *Agarivorans ablus* strain JK6, providing an ideal model for studying gene differentiation following duplication. Phylogenetic analysis revealed that these enzymes arose through gene duplication and subsequent divergence. Experimental assays demonstrated that, while they retained similar glycoside hydrolase activity, their agarolytic activity diverged significantly. We further explored their structural variations constrained by the protein’s 3D structural limitations, the development of specific localization linked to changes in enzymatic activity, and distinct expression patterns induced by different sugars. Notably, structural variations were primarily concentrated in the active site, while the overall backbone remained highly conserved. This study highlights gene differentiation following duplication as a key evolutionary strategy, facilitating the transition from single enzymes to complex functional systems.

## Main Text

Since Ohno’s seminal work in 1970^1^, gene duplication events have been recognized as key catalysts for functional innovation and biodiversity, playing a crucial role in augmenting genomic complexity and facilitating the emergence of novel gene functions^2-5^. This evolutionary mechanism allows duplicated genes to diverge through functional degradation, subfunctionalization, or neofunctionalization, thereby promoting adaptive evolution in response to environmental pressures^6-9^. Despite the acknowledged significance of gene duplication in evolutionary biology, numerous questions persist regarding how these genes attain functional differentiation and achieve regulatory refinement within specific ecological niches and intricate biological networks^10^.

To delve deeper into these mechanisms, microorganisms—particularly bacteria—serve as ideal models due to their remarkable adaptability and diversity^11^. They evolve complex enzymatic systems for specialized substrates via gene duplication and subsequent differentiation^12,13^. enabling them to navigate environmental shifts and competitive pressures. Among these enzymes, agarases—a crucial class of glycoside hydrolases that break down agar polysaccharides in seaweed^14,15^ —play an instrumental role in polysaccharide metabolism and the carbon cycle in marine bacteria^16^. Investigating these enzymes not only sheds light on the mechanisms of evolutionary and functional diversification but also enriches our understanding of microbial metabolic optimization and survival strategies in distinct ecosystems. However, in-depth studies on the functional differentiation of agarases following gene duplication, as well as their regulatory mechanisms in specific microbes, remain limited.

To address this gap, we conducted an in-depth investigation of the JK6 strain (*Agarivorans ablus*) from a unique deep-sea environment, which specifically degrades agar polysaccharides to obtain carbon nutrients.. By comparing the activity differences, cellular localization diversification, and expression pattern changes of four agarase proteins (PL3506, PL3511, PL3512, and PL3518), we elucidated their evolution and functional differentiation. Utilizing AlphaFold for highly accurate protein structure predictions, we examined structural changes and constraints during the differentiation process and achieved efficient and accurate predictions and modifications of protein activities. Through a comprehensive analysis of the agarase system in the JK6 strain, we demonstrated how simple adaptive processes can evolve into complex networks of ecological interactions^17^, revealing potential pathways for functional diversification following gene duplication. Moreover, our findings uncover bacterial adaptation mechanisms to environmental stresses through gene regulation and enzymatic system refinement, offering new insights into the role of microbes in the global carbon cycle and advancing the development of biotechnological applications utilizing agarases and other polysaccharide-degrading enzymes.

### Physiological Characterization and Genome Annotation of Strain JK6

The strain designated JK6, which was isolated in the present investigation, originates from a deep-sea water sample collected in the vicinity of the Mariana Trench. On L1 agar plates, the JK6 strain exhibited significant hydrolytic activity, forming clear halos indicative of agar degradation (Fig. S1). JK6 was identified as a strain of *Agarivorans ablus*. It has a genome size of 4.7 Mb and contains a 4,463 bp plasmid. Annotation identified 4,322 coding genes, 1,152 enzymes, and 280 transport proteins, which mapped to 282 known metabolic pathways. Positive selection analysis of single-copy orthologous genes revealed 53 genes under positive selection (Data S1), mainly involved in functions such as flagellar movement and ion transport, consistent with characteristics reported for deep-sea bacteria^18^.

### Functional Diversification and Evolutionary Analysis of GH50 Agarases in JK6 Strain

We utilized OrthoFinder to analyze the complete protein sequence files of more than 600 bacterial strains (Data S2), identifying a set of four agarases belonging to the GH50 family^14^, which were classified in the same Orthogroups. Through phylogenetic tree analysis of GH50 family proteins and structural similarity clustering of proteins containing the major glycoside hydrolase domains, we confirmed their evolutionary relationships (Fig. 1a). To establish the relationships among these four proteins, the distance agarase PL2878 was used as a common reference point.

**Fig. 1.**
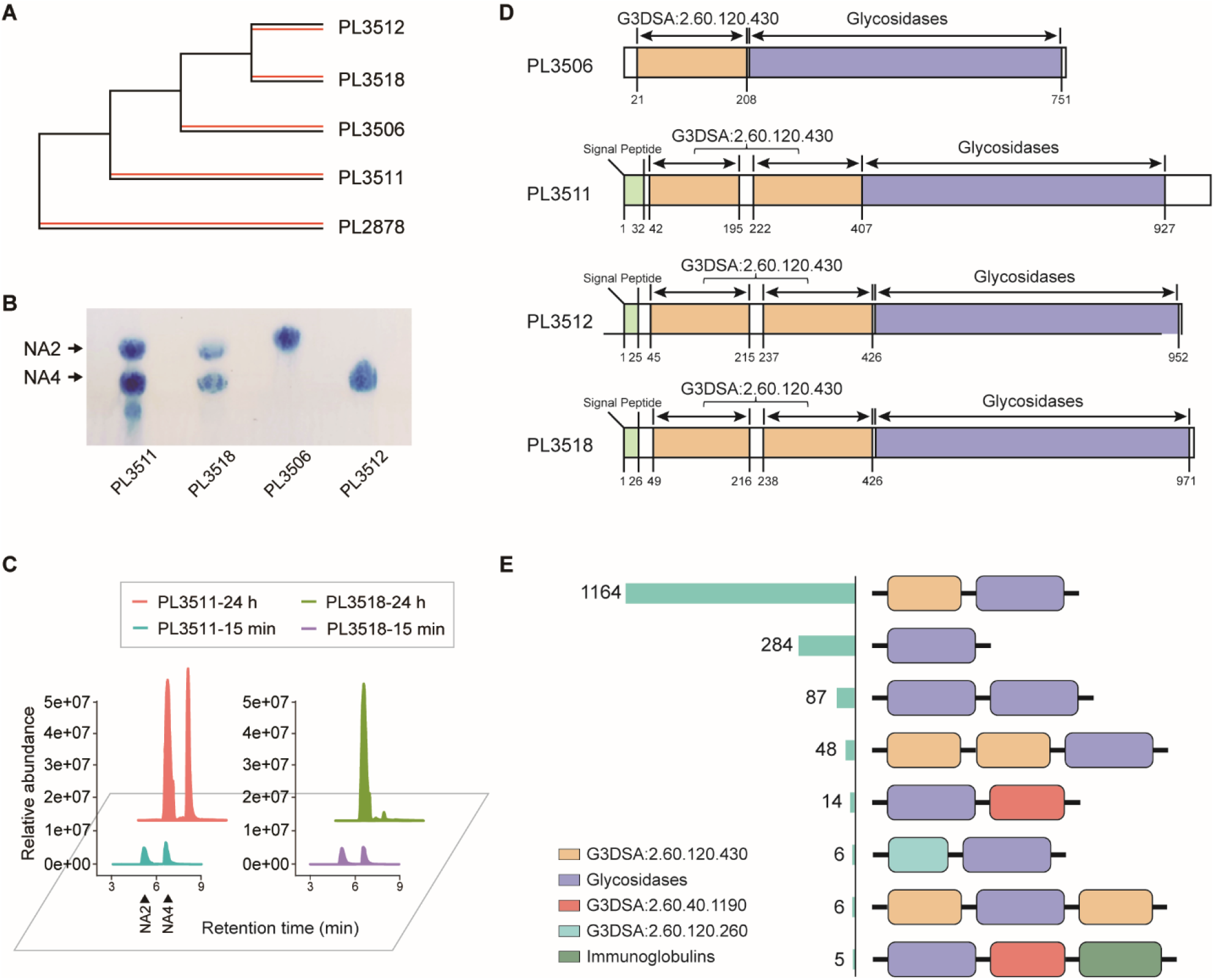
Gene Structure, Evolutionary Relationships, and Enzymatic Activity of Four Amylases. **(A)** Evolutionary topological relationships of four amylases and outgroup agarases, supported by phylogenetic trees based on protein sequences and clustering based on structural similarity. **(B)** The thin layer chromatography (TLC) results of the hydrolysis products obtained by treating excess agarose for 15 minutes with the four agarases expressed and purified from Escherichia coli (E. coli), which originated from the JK6 strain, using a mobile phase consisting of 1-butanol/acetic acid/distilled water (2:1:1, v/v/v). **(C)** The mass spectrometry abundance of the main polysaccharide products after 15 minutes and 24 hours of reaction with excess agarose for proteins PL3511 and PL3518 shows that at 15 minutes, the main product content between PL3511 and PL3518 is not significantly different. However, after 24 hours, most of the PL3518 products have been converted to NA2, while PL3511 still has a considerable proportion of NA4 accumulation, indicating a time-dependent difference in the hydrolytic activity between the two enzymes. **(D)** Gene3D annotation of the four proteins’ structural domains, with the G3DSA:2.60.120.430 domain marked as a CBM-like (carbohydrate-binding module-like) domain in Pfam annotations. **(E)** Structural domain composition types and corresponding quantities of GH50 family proteins in the CAZy (carbohydrate-active enzymes) database.

Although gene family expansion in prokaryotes is often considered to be driven by horizontal gene transfer^19^, our study ruled out the possibility of the four agarases. We speculate that these proteins were retained after multiple duplications from a common ancestor and were distributed within a gene cluster spanning approximately 23 kb (Fig. S4). Specifically, the genes *PL3511* and *PL3512* retained a tandem duplication pattern adjacent to each other. Considering that tandem duplications are usually highly unstable and without the action of natural selection, amplified gene arrays rapidly disappear from populations^2,20,21^. Therefore, natural selection may have occurred during the fixation process of the genes *PL3511* and *PL3512*.

After analyzing the complete protein sequences of 628 bacterial strains, we identified variations in the number of homologs for four proteins across different clades, reflecting their evolutionary relationships (Fig. S3). Homologous genes of the proteins PL3506 and PL3512 are primarily distributed across different clades and often do not coexist within the same species. Based on the inferred topology of the four-gene divergence, we observed that the earliest diverged PL3511 has the fewest homologs present among species, while the transitional PL3518 has slightly more homologs but still significantly fewer than PL3506 and PL3512. Additionally, we speculate that some degree of gene loss may have occurred during this process. In the genus *Vibrio*, we found that certain strains may have acquired this gene cluster through horizontal gene transfer, thereby ruling out the possibility of independent origins due to convergent evolution.

By analyzing the degradation products of agarose by these four agarases, we found that they have undergone significant functional differentiation (Fig. 1B). Specifically, PL3506 primarily produces NA2 as its main product, while PL3512 mainly generates NA4. Within a certain reaction time, PL3511 can produce NA2, NA4, and small amounts of NA6/NA8, whereas PL3518 mainly yields NA2 and NA4. Over time, PL3518 converts its main products to NA2 more rapidly than PL3511. Moreover, when testing the cleavage efficiency against the NA4 substrate, PL3518 indeed degrades it completely into NA2 more quickly than PL3511(Fig. 1C, Fig. 2C).

**Fig. 2.**
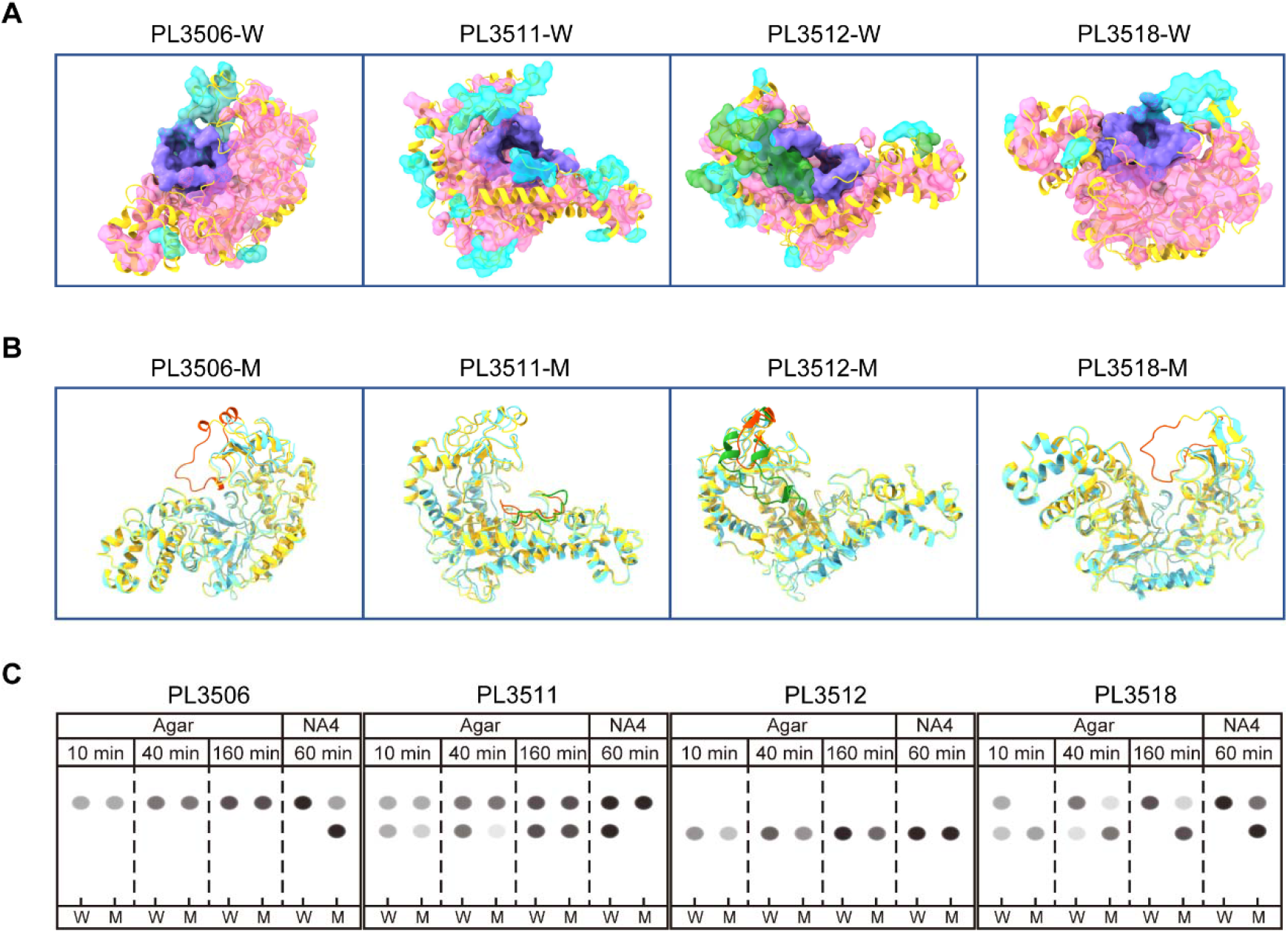
Structural Comparison of the Four Proteins and Changes in Enzymatic Activity After Directed Modification. **(A)** Differences (light blue), conserved regions (pink), and active sites (purple) in the structural comparison of each enzyme with the other three enzymes; **(B)** structural regions of directly modified enzymes (modification sites marked in red); **(C)** changes in hydrolytic activity toward two substrates for the four enzymes and their directed mutants, with “W” representing the wild-type enzyme and “M” denoting the modified mutant. The grayscale spots in the image represent hydrolysis products, with the depth of grayscale indicating the concentration of the products. In PL3506, which produces NA2 as a product, the loop differing from that in PL3512, which produces NA4, was removed. According to previous analyses of GH50 family protein structures, this loop structure is highly variable in sequence, and its presence or absence does not clearly correlate with whether the enzyme produces NA2 or NA4 as hydrolysis products. After modification, the percentage of NA4 cleaved by PL3506 decreased significantly, but the amount of agarose cleavage products produced remained unchanged, and only NA2 was produced, as predicted. Similarly, after removing the excess loop in PL3518, its rate of cutting NA4 also significantly decreased, and more NA4 accumulated when cleaving agarose, suggesting that the loop structure may only affect the binding or release of polysaccharides with agarase, thus influencing enzyme activity against different products. Additionally, experiments in which the CBM domain in PL3506 was removed revealed a significant reduction in the hydrolysis rate of NA4 and a change in agarose hydrolysis products, indicating that the CBM domain adjacent to the GH domain plays an important role in supporting the active center and enhancing polysaccharide binding.

Regarding gene structure, although there are discrepancies between the annotations from Gene3D and Pfam, the AlphaFold-predicted structures of the four proteins suggest that Gene3D’s predictions are more accurate^22^ (Fig. S2). Therefore, we selected the Gene3D results for subsequent expression validation of the glycoside hydrolase domains (Movie S1). All four proteins contain a glycoside hydrolase domain at the C-terminus. Except for PL3506, which retains a single CBM-like domain, the other three proteins possess two CBM domains located at the N-terminus^23^. Structural comparisons of the CBM domains and glycoside hydrolase domains among these proteins revealed a high degree of similarity (Movie S2). According to annotation statistics of GH50 family proteins, the combination of two CBM domains plus one glycoside hydrolase domain is relatively rare in the GH50 family, which may be associated with the oligotrophic ecological environment in which the strain resides^24^.

In summary, our comprehensive analysis of the four GH50 agarases in the JK6 strain reveals a clear evolutionary trajectory and functional diversification resulting from gene duplication events. Phylogenetic and structural analyses indicate that these agarases originated from multiple duplications of a common ancestor and have since diverged in both sequence and function. PL3511 appears to retain the most ancestral enzymatic activity, producing a variety of degradation products and serving as a link to the original enzyme function. PL3506 and PL3512 have undergone subfunctionalization, specializing in the production of NA2 and NA4, respectively, while PL3518 represents an intermediate transitional state. The distribution of homologous genes across different bacterial clades and the presence of tandem duplications suggest that natural selection has acted to retain and specialize these genes in response to environmental pressures, particularly in oligotrophic deep-sea conditions. These findings highlight the role of natural selection in shaping the functional diversity of duplicated genes and provide valuable insights into the mechanisms of bacterial adaptation and evolution within specific ecological niches.

### Modifications in Activity within the Constraints of Protein Structural

The interaction between structure and function in the evolution of proteins exhibits a high degree of complexity^25,26^. Typically, the conservation of amino acid sequences is crucial for maintaining the specific three-dimensional structure of a protein, which plays a key role in its biological function^27-30^. As a result, although mutations may occur in the evolutionary process, conservation at key positions ensures the stability of protein structure and function. This delicate equilibrium reveals that sequence variation drives protein evolution while being constrained by structural and functional limitations^31^, reflecting the complexity of biomolecular adaptability and evolutionary dynamics.

In this study, we used AlphaFold to predict the structures of our target proteins, revealing that all four agarases displayed mixed folding characteristics. At the enzyme’s N-terminus, they included one to two β-sandwich domains (CBM-like domains), while the C-terminus exhibited a sophisticated (α/β)8 barrel (glycoside hydrolase domain). Two CBM domains were connected by a linker region of approximately 22 amino acids, with the CBM domain adjacent to the glycoside hydrolase domain located near the (α/β)8 barrel (Fig.1D).

Using the domain ranges defined by Gene3D, we extracted the glycoside hydrolase domains and performed AutoDock Vina docking experiments using polysaccharide molecules from NA2 to NA8 as substrates. After adjusting parameters and repeating the experiments multiple times, we identified amino acids within 5□Å of the sugar molecules, considering these residues as regions related to the active center (Fig.2A/B). We then conducted pairwise comparisons of the extracted glycoside hydrolase domains using USalign, employing the spatial vector modulus of the TM-score and RMSD between two proteins as the similarity score Vd, and constructed a matrix to build a clustering tree. The results were largely consistent with the branching relationships in the gene phylogenetic tree (Fig.1A, Fig.S6), providing valuable reference information. It is important to note that due to potential alignment differences caused by the flexible structures of multi-domain proteins, this study focused solely on structural similarity analysis of the core domains. Based on structural similarity clustering, the four agarases were distinctly divided into four groups.

Interestingly, unlike the commonly observed mutations that occur mainly in non-active sites of enzymes^28^, we found that the primary differences among the four agarases are located at their active centers, while the overall protein scaffolds remain highly conserved. As shown in Table S1, pairwise comparisons of the glycoside hydrolase domains revealed RMSD values less than 1.9□Å and average TM-scores higher than 0.9. However, this structural conservation contrasts sharply with the amino acid sequence similarities, which are below 50%. Even when the comparison is extended to proteins from Vibrio species strains of different genera, a similar pattern is observed. The calculated Ka/Ks values are much less than 0.5, indicating that the overall sequences are under significant purifying selection.

Protein evolution is influenced by multiple factors, including the structure of the genetic code and the conservation of amino acid physicochemical properties. To further characterize structural conservation, we calculated the ratio of amino acid residues with pairwise structural differences greater than 1□Å to those with differences less than this threshold. This ratio is also much less than 1, confirming the high level of structural conservation despite sequence divergence.

While the homologous proteins exhibit similarity in their main structures, their local structural differences, particularly at the active centers, are noteworthy. Considering that the Ka/Ks value reflects selection pressure on proteins at a global level, we found that large regions of the proteins are under purifying selection to maintain overall structure and stability. However, this overlooks the fact that only a few sites are under positive selection. Using a sliding window of 120 codons, we examined the regional variations in the Ka/Ks values and found that the enzyme active center regions indeed fall within sequences under positive selection.

Our analysis indicates that although significant divergent variations have occurred in both DNA and protein sequences, these mutations have been constrained by protein structural requirements. This constraint has led to divergent mutations that provide adaptive advantages specifically at the active centers, driving functional diversification while preserving overall structural integrity.

To investigate the functional implications of these structural differences at the active centers, we conducted site-directed mutagenesis experiments (see Fig.2). In PL3506, which produces NA2 as its product, we removed a differential loop that distinguishes it from PL3512, which produces NA4. Structural analysis of GH50 family proteins suggests that this loop exhibits high sequence diversity and is entirely absent in many cases; moreover, its presence or absence is not clearly correlated with whether the enzyme produces neoagarobiose or neoagarotetraose as hydrolysis products. The modified PL3506 showed a significant decrease in the rate of NA4 cleavage but did not change the type of products from agarose hydrolysis, still producing only NA2, consistent with our predictions.

Similarly, after removing the redundant loop in PL3518, its rate of NA4 cleavage decreased significantly, and more NA4 accumulated during agarose hydrolysis. This suggests that the loop structure may affect only the binding or release of polysaccharides by the agarase, thereby influencing enzyme activity toward different products. Additionally, we performed deletion experiments on the CBM domain in PL3506 and found that the modified PL3506g exhibited a significantly reduced hydrolysis rate of NA4, and its hydrolysis products from agarose changed. This indicates that the CBM domain adjacent to the glycoside hydrolase domain plays an important role in supporting the active center and enhancing polysaccharide binding.

We also experimented with fully conserved amino acid sites in the glycoside hydrolase domain. Mutating the Glu residue adjacent to the active center in PL3506 to Gly resulted in changes in protein activity, indicating that these conserved sites play critical roles in protein function.

Furthermore, to validate our findings across different species, we performed synthetic expression experiments on the four homologous proteins from a *Vibrio astriarenae* strain (HN897) of different genera^32^. These genes can be ruled out as convergent results from independent evolution, providing appropriate controls for horizontal comparison. The experimental results showed that the homologous protein of PL3506 in VSP strains has NA2 hydrolysis activity and exhibits monosaccharide hydrolysis activity during agarose degradation without accumulating large amounts of NA2. Additionally, the homologous proteins of PL3511, PL3512, and PL3518 also produced certain monosaccharide products, but these products were not generated by hydrolyzing NA2, suggesting that they may cleave non-NA2-type β1-4 glycosidic bonds at the outermost ends of agarose.

In conclusion, these findings highlight that structural divergence at the active centers, constrained by the overall protein structure, drives the functional diversification of agarases following gene duplication. This provides valuable insights into the mechanisms of enzyme evolution and bacterial adaptation, emphasizing the role of targeted mutations in active sites in developing new enzymatic functions while preserving structural stability.

### Changes in Subcellular protein distribution

Retention of duplicated proteins offers significant biological advantages, particularly in the subcellular localization differentiation of proteins, which facilitates more precise regulatory expression and operational efficiency^33^. Some studies have indicated that functional differentiation caused by subcellular localization is crucial to the biological function of proteins. In this study, we inferred the subcellular localization of the four agarases through protein localization prediction, analysis of intracellular and extracellular proteomic abundance under different culture conditions, and expression localization of complete proteins in Escherichia coli (see Fig.3).

**Fig. 3.**
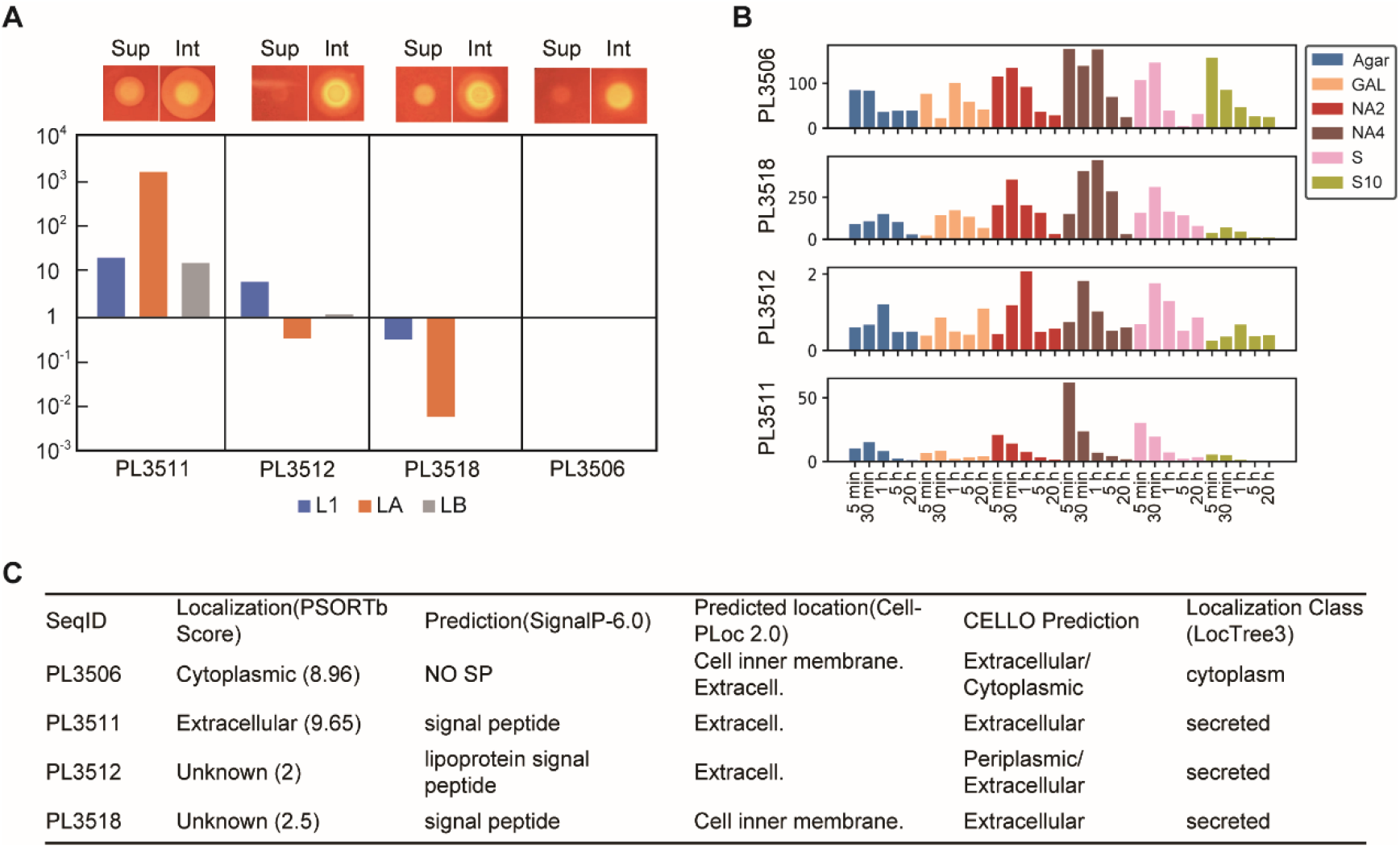
Changes in the Localization and Expression of the Four Proteins. **(A)** The top section shows the differences in agarolytic activity on agar plates supplemented with concentrated proteins (Sup) and cell lysates (Int) during the expression of the four enzymes in *Escherichia coli*. The bottom section compares the mass spectrometry ratios of the supernatant to intracellular proteins under three different culture conditions (L1 medium, L1A medium, and L1B medium). No mass spectrometry was detected in the supernatant for 3518 under L1B culture or for 3506 under any culture conditions; hence, 3518 was used as a blank. Notably, the ratio of intracellular to extracellular accumulation significantly changed under different culture conditions. Especially under agarose induction, PL3511 and PL3518 showed greater and lower proportions of extracellular secretions, respectively. **(B)** qPCR results of the four proteins cultured under five different sugar conditions (agar-L1 medium with 5% agar blocks, GAL-L1 medium with 0.1 g/L galactose, NA2-L1 medium with 0.1 g/L NA2, NA4-L1 medium with 0.1 g/L NA4, S-L1 medium with 0.1 g/L agar oligosaccharides, and S10-L1 medium with 1 g/L agar oligosaccharides). **(C)** Localization predictions for the four proteins using different software.

Our results showed that PL3511 is an extracellular secreted protein, while PL3506 is mainly localized intracellularly. PL3512 behaves as a periplasmic protein, whereas PL3518, although exhibiting extracellular secretion characteristics, accumulates more significantly inside the cell. Notably, the proportions of intracellular and extracellular accumulation exhibit significant changes under different culture conditions. Specifically, under agarose induction, PL3511 and PL3518 show higher and lower proportions of extracellular secretion, respectively. This finding is closely related to the functional differentiation of the four agarases’ activities, suggesting that cells can degrade agar-containing polysaccharides into longer chain polysaccharides extracellularly. These polysaccharides are then transported into the periplasm, where they are further hydrolyzed into NA4 by PL3512, and finally degraded into NA2 by intracellular PL3506, before being metabolized through subsequent monosaccharide hydrolysis and utilization pathways.

The four agarases exhibit differences in subcellular localization and secretion properties. This functional and localization differentiation not only enhances the cells’ efficiency in utilizing agarose but also exemplifies the biological advantage of retaining duplicated proteins, achieving precise regulation through diversification of subcellular localization.

### Ancestral sequence reconstruction analysis

Reconstructing ancestral enzymes to explore the evolutionary mechanisms and fates of duplicated genes is a viable strategy^34,35^. Our study reconstructed the ancestral sequences of four agarases based on the GH50 gene family tree and analyzed their evolutionary history by comparing the three topological structures (Fig. S7). Examination of subcellular localization predictions for ancestral protein sequences at various nodes revealed that ancestors related to PL3511, PL3512, and PL3518 largely retained the same localization as their contemporary counterparts, whereas the ancestors of PL3506 exhibited distinct localization patterns (Data S6). Our analysis suggested that subcellular adaptability is a key mechanism for the functional diversification of duplicated genes, as evidenced by the absence of intracellular localization predictions for PL3506 ancestors, indicating that functional diversification may precede localization differentiation, and indicating that the evolution of localization and function might not always be synchronous.

### Changes in Gene Expression Regulation

In the process of functional differentiation following protein duplication, the regulation of gene expression is a key mechanism^2,36^. Our study revealed distinct responses and expression patterns of these four agarase genes in different sugar environments. This differential expression suggests that bacteria employ a more efficient protein hydrolysis method to acquire nutrients under limited resources, thereby gaining a competitive advantage in terms of survival.

We employed qPCR technology to assess the expression of these four proteins under various sugar concentrations (Fig. 3B). The results demonstrated that these proteins exhibit distinct expression patterns in response to different sugars or sugar concentrations, especially under the induction of NA2, NA4, and oligosaccharides. The PL3511 protein quickly reached a transcriptional peak, while PL3512 and PL3518 showed differences in response to NA2 and NA4. Specifically, PL3512 rapidly reached its peak under NA4 induction but was slower in NA2; conversely, PL3518 responded quickly under NA2 induction and relatively more slowly under NA4, indicating that in the initial induction, the corresponding product of the enzyme has a negative feedback function to induce enzyme transcription. Additionally, PL3506 exhibited the fastest transcriptional response under relatively high concentrations of oligosaccharides.

These results indicate that the four agarase genes are specifically regulated in response to different sugar environments, suggesting that bacteria can adapt to varying nutrient conditions by modulating enzyme expression. Additionally, the presence of a negative feedback mechanism highlights the fine-tuned regulation between gene expression and metabolic products, enhancing bacterial competitiveness in resource-limited environments.

### Covariation Evolutionary Rate Analysis

To study the coevolutionary history between genes, we calculated the coevolutionary rates^37^ of gene pairs to reveal their covariation characteristics (Data S7, Fig. S8). Within the set threshold range, we found that *PL3511* did not exhibit a significant correlation with the other three agarase genes (*PL3506, PL3512*, and *PL3518*). This may indicate that PL3511 has greater functional and evolutionary independence from the other three enzymes. Considering the evolutionary rates^38^ and the average pairwise identity^39^ between sequences, there were significant differences between PL3511, PL3512, PL3518, and PL3506. The evolutionary rates from low to high were PL3506, PL3518, PL3512, and PL3511, while the average pairwise identity mean values from high to low were PL3512, PL3518, PL3506, and PL3511. These results suggest that PL3511 differentiated earliest from the other three agarase genes, with PL3518 possibly in an intermediate transitional state between PL3506 and PL3512. Although the other differentiation processes among these genes are not entirely clear, it can be inferred that they have undergone complex pathways of functional and evolutionary differentiation. This not only reveals their unique roles in metabolic and physiological processes but also provides important clues for further understanding their functions in specific cellular environments.

## Discussion

### Divergence Patterns

The most common fate of duplicated genes is the deletion of some members, returning the gene to a single-copy state^8,40^. In addition, duplicated genes may undergo nonfunctionalization (to maintain an appropriate level of function), subfunctionalization (functional partitioning), or neofunctionalization (functional diversification)^8,41^. Since Ohno’s proposal, several more detailed models have been accepted, such as duplication-degeneration-complementation (DDC)^3^, escape from adaptive conflict (EAC)^42^, and innovation-amplification-divergence (IAD)^43^. However, overall, the fate of duplicated genes and their impact depend to some extent on the timing of the duplication event. Here, we will briefly compare the divergence of these four proteins following duplication, focusing on aspects such as their evolutionary history, structural differences, functional changes, and expression patterns, to determine which model of divergence they most closely align with.

By analyzing the different types of these four agarase proteins, we observed that the activities of PL3506 and PL3512 were markedly different from that of the more primitive version, PL3511. Product specificity suggests subfunctionalization, while enzyme mechanisms imply neofunctionalization. Therefore, we only confirmed that functional differentiation occurred without making overly detailed distinctions. Our analysis indicated that PL3511 is more closely related to the ancestral enzyme, hinting at preexisting multifunctionality. PL3506 and PL3512 specialize in enzymes with single subfunctions, which can resolve conflicts between different transcriptional demands. Additionally, protein localization differentiation likely occurred after enzymatic activity differentiation, raising questions about the cause of enzyme diversity. We propose that differentiation aligns more with the EAC model, suggesting that adaptation and localization changes postduplication. Enzyme evolution is complex, and understanding it fully requires considering factors such as metabolic network integration. Our current models (Fig. 4) are preliminary and will evolve with more data.

**Fig. 4.**
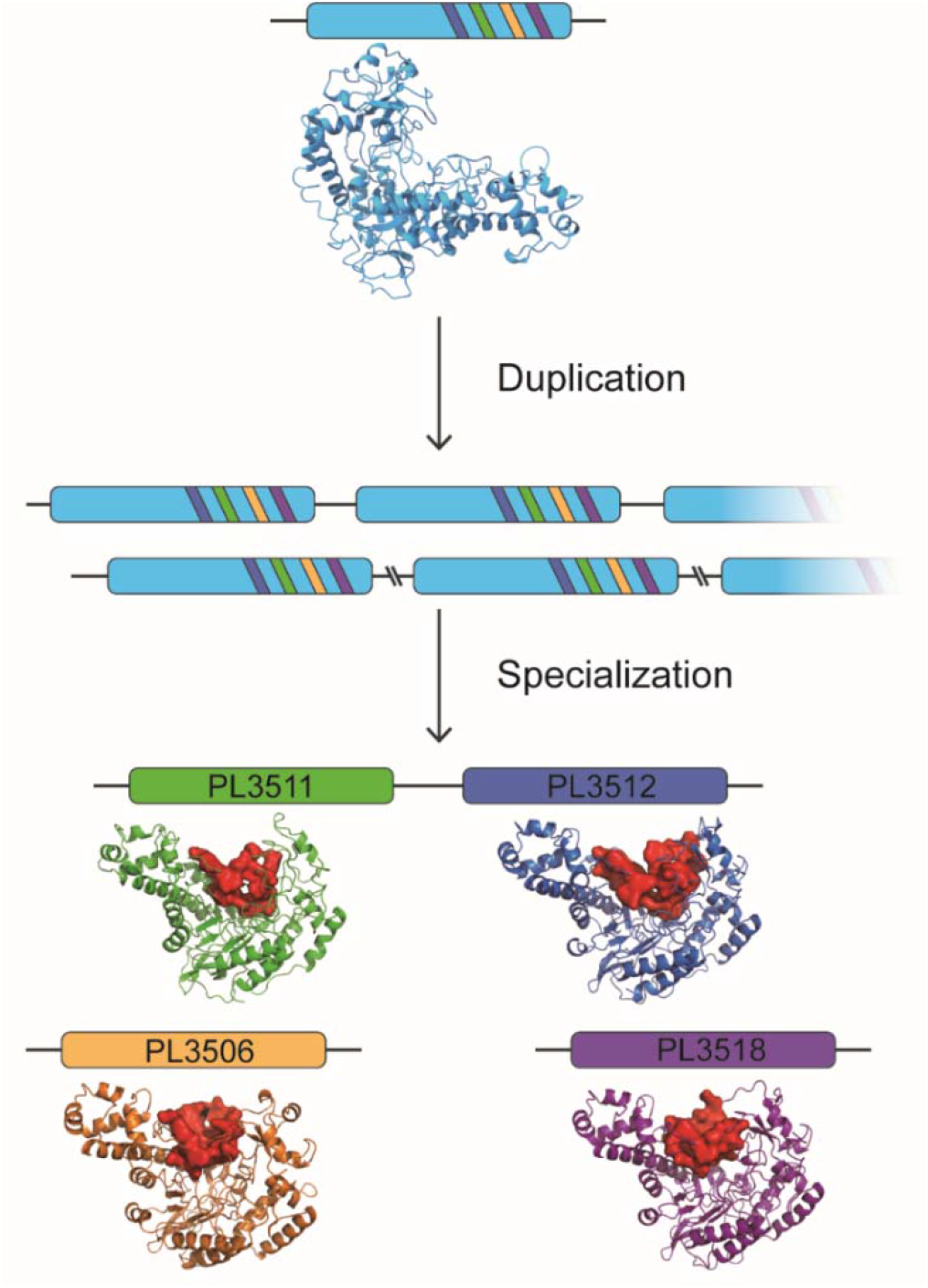
Protein Duplication and Divergence Model. An ancestral gene with multifunctional activity may be duplicated and retained under selective pressures such as dosage requirements, and subsequently, each duplicated gene copy diverges through mutations. From the perspective of the final products, some genes tend toward specialization, leading to the resolution of the conflict between the transcriptional regulation of the two functions and gaining advantages in the precise regulation of enzyme activity and expression levels, thus gradually forming an enzyme control network for hydrolyzing the substrate. During the process of mutation, due to the similarity in enzyme functions, constraints are observed in the protein’s 3D structure, whereby although there are significant changes in sequence similarity, the main structure of the protein remains consistent, while major variations in structure occur at the enrichment of the enzyme active center. As we explore further into the intricacies of enzyme evolution, particularly within the GH50 family of glycoside hydrolases, we found that they maintain fundamentally consistent spatial structure similarity, showing only local structural differences. This pattern is akin to bird nests built on different trees in a forest: while the basic form of the trees is consistent, the position and type of nests vary among trees due to differences in birds. This may be because highly specific enzymes need to function effectively in the presence of structurally diverse polysaccharide substrates, and given the relatively small variety of basic units and linkages in polysaccharides, different glycoside hydrolases need only slight adjustments to address various polysaccharide structures.

### Integrated Agar Degradation Pathway Mediated by Diverse Agarases in JK6 Strain

Bacterial communities are typically not composed of single cells but rather possess a certain structure, such as biofilms^44^. Against this backdrop of complexity, including constantly changing environments and heterogeneity among cells and/or species, bacteria have evolved complex enzyme systems for selective expression, thus addressing environmental challenges in an economical and efficient manner. In the JK6 strain, in addition to this group of homologous agarases, we also identified several agarases with different localizations and hydrolytic activities through genome annotation and similarity comparisons of known agarases based on AlphaFold-predicted structures. For example, PL3515^45^ is involved in the final step of agar metabolism and has exo-acting α-1,3-agarase activity in AHG (3,6-anhydro-α-L-galactose) and GAL (D-β-galactose), and PL2878^46^ is predicted to be located on flagella.

In natural environments, bacteria coexist with other bacteria and organisms, engaging in a variety of interactions^47^. Based on the results of previous studies^47,48^, we speculated that the metabolic pathway of the JK6 strain could be determined via agar (Fig. S9). Initially, agar in the environment is hydrolyzed into longer polysaccharide units by exoenzymes and flagellar-located agarases.

These two protein types may produce different induction signals for free and fixed agar in the environment (the intracellular cyclic di-GMP phosphodiesterase (*PL3852*) increases with the accumulation of agar signals, inducing the transition from motility to attachment and the formation of biofilms in the strain), leading to distinct responses. Upon receiving short-chain sugar signals, the cell synthesizes and secretes large quantities of PL3518 and PL3511, combined with cell motility, to further breakdown fixed agar into short-chain polysaccharides. Together with polysaccharide transport proteins, shorter chain polysaccharides are transported to the periplasm, where they are hydrolyzed into NA4-sized units by enzymes located there, such as PL3512, and then transported into the cell via polysaccharide ABC transporters. Intracellular PL3506 hydrolyzes it into disaccharides, which are then hydrolyzed into monosaccharides by PL3515, which subsequently enter metabolic pathways such as AHG metabolism (PL0686, PL3528, and PL3525) and the Leloir pathway (PL1674, PL1675, and PL1676).

Notably, the agarase system identified in the JK6 strain is not organized into a typical operon system, and these enzymes are relatively dispersed throughout the genome. Although PL3506, PL3511, PL3512, and PL3518 are clustered together, no representative complete operon elements are found nearby. While two-component systems were identified, no complete sucC/D transport system was detected, suggesting the possibility of alternative transport pathways. Given the relatively nutrient-poor marine environment, bacteria are expected to possess regulatory systems that respond to fluctuations in nutrient availability. Based on the ecological model of agar degradation in the JK6 strain, it is likely that other unconventional regulatory systems exist, which warrant further exploration.

## Conclusions

Our study systematically explored the evolution and functional differentiation of the agarase system in the JK6 strain. By analyzing the structural and functional characteristics of four agarases, we revealed that, following gene duplication, these enzymes were subject to strict structural constraints, with functional differentiation primarily occurring through changes in the active sites. The close relationship between subcellular localization and functional divergence, along with distinct expression patterns in various nutrient environments, suggests that these enzymes adapt to complex environmental changes through selective expression. Additionally, the dispersed distribution of the agarase system across the genome indicates the potential involvement of non-traditional regulatory mechanisms. This research provides new insights into microbial adaptive evolution and gene regulatory networks, laying a foundation for the biotechnological application of agarases and other polysaccharide-degrading enzymes.

## Supporting information

data1-9 and mov1-3

## Acknowledgments

We thank Prof. Weicheng Cui and his technical team members of the 2020 joint cruise to the Mariana Trench of Westlake University and Shanghai Ocean University. We appreciate Dr. Xiaobo Li, Dr. Ying Zhen and Dr. Yanlei Feng for their advice and assistance in writing the article. We thank Dr. Mingzhu Fan, Xiaoyan Xu and Jinheng Pan from the Biomedical Research Core Facilities at Westlake University for assistance and discussion during the experiment. We thank the Westlake University HPC Center for computational support. We thank Lei Gao, Yingping Gao and Fang Xiao at the Westlake University Microscopy Core Facility for advice and assistance in microscopy sample preparation and data collection. We thank Huican Li from Westlake University for his help in the cultivation and collection of *Escherichia coli* and all colleagues on the outbound team, especially Xinyu Huang, for their help in providing the experimental materials needed for sample collection.

## Funding

Zhejiang Provincial Natural Science Foundation of China LQ20C040001 (KLZ) National Natural Science Foundation of China No. 32100484 (KLZ)

## Author contributions

Conceptualization: KNG

Methodology: KNG, KLZ, YYQ

Investigation: KNG, YYQ

Visualization: YYQ, TTY, KNG

Funding acquisition: KLZ

Project administration: KLZ, KNG

Supervision: KLZ

Writing – original draft: KNG

Writing – review & editing: KNG, YYQ, TTY, KLZ

KNG and YYQ made equal contributions to this work.

## Competing interests

The authors declare that they have no competing interests.

## Data and materials availability

The bacterial strain JK6 used in this study will be shared upon request. The specific code and Alphafold-predicted structures mentioned in the article are available on the GitHub link (https://github.com/jkgkn/JK6). The raw sequencing data can be obtained from SRR27869121, SRR27869122, and SRR27869123. Upon request, the corresponding author can provide any further information needed for the reanalysis of the data presented in this paper.

## Notes

### Competing Interest Statement

The authors have declared no competing interest.

### Summary of Updates

This version has revised the abstract, enriched the content in certain paragraphs of the main text, and updated the content in B and C of Figure 1.

