## Supplementary material for "Functional Diversification of Gene Duplicates under the Constraint of Protein Structure": data1-9 and mov1-3

### Materials and Methods

Abbreviations:
D-Gal, D-β-galactose;
L-AHG, 3,6-anhydro-α-L-galactose;
NAOS, neoagarooligosaccharide;
DP, degree of polymerization;
NABH, α-neoagarobiose hydrolase;
GH, glycoside hydrolase;
NA2, neoarabinose;
NA4, neoagarotetraose;
NA6, neoagarohexaose;
NA8, neoagarooctaose;
A3, agarotriose.

All reagents and chemicals were purchased from Sigma–Aldrich (St. Louis, MO) unless otherwise specified.

#### Field Sample Collection, Laboratory Isolation and Identification, and Microscopic Observation

During the 2020 joint cruise to the Mariana Trench of Westlake University and Shanghai Ocean University, water samples were collected at depths of 0-4 km at various stops along the route, and sediment samples were collected at a depth of 6 km at coordinates 18.5047 and 132.0094. Microalgae from the deep sea were enriched and isolated by spreading 100 µL of 1-6 km samples (water samples were centrifuged for 10 minutes at 5000 g, the supernatant was removed, and 100 µL of the bottom liquid was retained; sediment samples were precipitated with sterilized L1 culture medium, and the supernatant was collected) onto L1 culture plates. Unexpectedly, colonies that produced agar-hydrolyzing pits were observed and purified by streaking on fresh plates multiple times. Ultimately, strain JK6, which exhibited the largest hydrolysis zone in a unit of time, was selected for subsequent experiments. The strain was preserved long-term at -80°C in 5% glycerol and cultured in L1 media (see L1 media from NCMA) supplemented with 0.2% agar (L1A) or 1% LB broth (L1B) for liquid culture and 1.5% agar for solid culture.

A nearly full-length 16S rRNA gene sequence was PCR amplified using the universal primers 27F and 1492R to identify the strain[1,2]. The obtained 16S rRNA gene amplicon sequence was deposited in the National Center for Biotechnology Information (NCBI) database and annotated through Sanger sequencing using the National Center for Biotechnology Information (NCBI) Basic Local Alignment Search Tool (BLAST).

After Gram staining, brightfield images were captured using a Novel DN-107T microscope equipped with an Optronics digital camera. The sample preparation process for scanning electron microscopy (SEM) involved collecting the JK6 bacterial aggregates formed in liquid culture, fixing them in a fixation solution (pH=7.2) containing a mixture of 2% paraformaldehyde (PFA) and 2.5% glutaraldehyde (GA) for 1 hour at room temperature, rinsing the samples three times with 0.1 M PB (pH 7.2-7.4) buffer for 10 minutes each time at 4°C, fixing them with 1% osmium tetroxide (in 0.1 M PB) on ice for 1 hour, rinsing the samples three times with ddH2O for 15 minutes each time at room temperature, dehydrating them with 30%, 50%, 70%, 95%, and 100%×3 ethanol for 10 minutes at room temperature, collecting the samples in a paper bag, drying them using a critical-point dryer (Leica EM CPD300), and finally sputter-coating them 5 nm thick with Leica EM ACE600. Imaging was conducted using a scanning electron microscope (Zeiss Gemini SEM Crossbeam 550).

**Genome Sequencing Assembly and Preliminary Annotation**

The genomic DNA of strain JK6 was extracted using the TIANamp Bacteria DNA Kit and then split into two batches for sequencing using both Illumina next-generation sequencing (PE150, Beijing Novogene Bioinformatics Technology Co., Ltd.) and Oxford Nanopore (PromethION, by Wuhan Benagen Technology Co., Ltd.). All experimental procedures, including sample quality testing, library construction, library quality testing, and library sequencing, followed the standard protocol provided by the sequencer manufacturers.

The genome assembly process consisted of three steps: raw read filtering, coassembly of second- and third-generation sequencing data, and polishing of the assembly results. Raw short reads from Illumina sequencing were quality controlled using Fastp (-q 30 -u 10 -n 5 -l 130)[3], and raw long reads from Nanopore sequencing were filtered using the clean reads provided by the company (removing sequences with an average quality value of less than or equal to 7). The clean short reads and long reads were then coassembled using SPAdes-3.15.2-Linux[4], wtdbg2[5], minimap2[6], SAMtools[7] and Unicycler[8] to generate complete genome sequences. The assembly results of each version were evaluated using Quast.py and MUMmer[9] to examine the assembly statistics and collinearity between different versions. The best version was polished twice using pilon-1.24.jar[10]. Gene annotation software was used to compare the annotation results before and after polishing. As there was almost no difference between the two results, the polished genome version was selected as the complete genome sequence. To annotate the selected reference genome, annotation results from different software programs, including GeneMarkS[11], Prodigal[12] and Prokka[13], were compared based on the number of annotated genes and the consistency of the results obtained by different methods. Prokka was chosen as the final gene annotation file.

We collected datasets such as "UniProt-reviewed_yes" prior to February 25, 2022, and constructed indexes for them. The protein sequence of JK6 was aligned against each dataset using the diamond[14] BLASTP command, with parameter settings of --evalue 1e-5, --id 30, and --max-target-seqs 1, to annotate homologous proteins for each protein sequence.

**Horizontal gene transfer analysis**

We used BLASTP searches against the NCBI nonredundant protein database to identify genes that may have originated from HGT[15,16]. To ensure that the hits represented likely homologs, we retained only hits with high-scoring pairs accounting for at least 90% of the query length and an overall amino acid sequence similarity of at least 40%. Genes with the best hit that lacked a genus-level taxonomic assignment or were derived from metagenomics surveys were manually examined for more reliable inference. A gene was considered putatively acquired if more than half of the top five hits were from other genera.

**Comparative Genomics Analysis**

Comparative genomics analysis of *Agarivorans* species using Orthofinder[17] and Get_homologues[18]. In this study, we downloaded 629 genomes (Data S2) containing agarase protein sequences from NCBI using a dataset tool. The genomes were annotated using Prokka, and Orthofinder.py and get_homologues.pl were used to compute the faa and gbf files for all species. Based on the results, closely related *Agarivorans* species were selected for pangenomic analysis following the software instructions (partial strain information is provided in the attached Data S8). METABOLIC-C.pl[19] was used to annotate the *Agarivorans* genomes, and METABOLIC-G.pl[19] was used to annotate the protein sequences, followed by comparison of the similarities and differences among different genomes.

**Positive Selection Analysis and Ancestral Sequence Reconstruction by PAML**

We selected the genomes of *Agarivorans* strains collected from shallow seas as background controls (Data S8) and used OrthoFinder to identify orthologous genes in this group of strains compared to JK6. Only single-copy orthologs with full-length protein sequences longer than 0.7 times the longest sequence in all strains were retained. The corresponding DNA sequence files and tree files were filtered accordingly and used for subsequent selective pressure and convergence analyses.

The influence of natural selection on protein-coding genes was inferred by estimating the ratio of nonsynonymous to synonymous substitutions ($\omega$). In brief, $\omega > 1 = 1$ and $< 1$ indicate positive selection, neutral evolution and negative selection, respectively. We used the program codeml for selective pressure analyses in Phylogenetic Analysis by Maximum Likelihood (PAML)[20]. The p values were estimated assuming a null distribution that is a 50:50 mixture of a $\chi 2$ distribution and a point mass at zero. The p values were corrected for multiple testing, and a false discovery rate (FDR) cutoff of 0.05 was used.

Ancestral sequence reconstruction was performed using PAMLX[21], with the following parameters: noise = 4, verbose = 1, seqtype = 2, aaRatefile = wag.dat, model = 3, fix_alpha = 1, RateAncestor = 1, Small_Diff = 5e-6, fix_blength = 2, method = 0.

**AlphaFold Structure Prediction and Docking Comparison**

The genome proteins of JK6 were compared to those in the NCBI database, and those with 100% sequence similarity were downloaded from the AlphaFold Protein Structure Database (https://alphafold.ebi.ac.uk/). For the remaining 1000+ proteins, monomer structures were predicted using locally deployed AlphaFold[22,23], and the agarases were additionally predicted using RoseTTAFold[24] at https://robetta.bakerlab.org/. Polysaccharide ligands were downloaded from PDBe (Data S9), and full-genome protein docking was performed using TANKBind[25]. The docking of agarase proteins and candidate agarase proteins was carried out using Vina[26,27], with the docking box confirmed by AutoDockTools-1.5.7. Using the structural domain ranges defined by Gene3D, the glycoside hydrolase domains were extracted and subjected to AutoDock Vina docking experiments with polysaccharide molecules from NA2 to NA8 as substrates. After adjusting the parameters and repeating the experiments multiple times, amino acids within 5 Å of the sugar molecules were counted, with those exceeding a threshold considered part of the active center region. The docking of glycoside hydrolase domains and related proteins was accomplished in bulk using Python scripts (https://github.com/jkgkn/JK6).

Structural similarity comparisons between the whole-genome proteins and the obtained homologous glycoside hydrolase domains were conducted using US-align[28]. The magnitude of the spatial vector composed of TM1, TM2, and RMSD was utilized as a measure of distance to compare structural similarity between two proteins. The specific calculation formula is $\sqrt{(1 - Tm_1)^2 + (1 - Tm_2)^2 + (RmSD\ /\ 10)^2}$. USalign was used to pairwise compare the extracted glycoside hydrolase domains of the proteins, with the structural similarity score Vd represented by the spatial vector magnitude of the TM scores and RMSD score between two proteins, and a clustering tree was constructed based on this matrix. The results were generally consistent with the branching relationships of the gene phylogenetic trees, providing valuable insights. Notably, due to the potential for large alignment discrepancies caused by the loose structure of multidomain proteins, this study focused solely on structural similarity analysis of core domains. According to structural similarity clustering, the four agarases were clearly divided into four groups (Fig. S6).

**Calculation of Ka/Ks**

We utilized KaKs_Calculator3.0[29] to calculate the selective pressure. Specifically, we prepared pairwise comparison homolog group files (PLs.homologs), a file containing multiple amino acid sequences (prp.fasta), and a file containing multiple nucleotide sequences (cds.fasta). Then, we executed the ParaAT.pl script[30] with the parameters -h PLs.homologs -n cds.fasta -a prp.fasta to generate the AXT sequence files required for calculating Ka (nonsynonymous substitution rate) and Ks (synonymous substitution rate). For the genetic code, we selected the codes for bacteria, archaea, and plant plastids. Regarding the calculation method, we chose to use all substitution models (ALLs) for the computations. Although homologous proteins exhibited overall structural similarity, their significant functional differences indicated the presence of variations. Considering that Ka/Ks values globally reflect selection pressure on proteins, there are extensive protein regions under purifying selection to maintain the overall structure and stability of the enzyme, but this overlooks the fact that only a few sites are under positive selection. Hence, we examined the interval changes in the ka/ks values of proteins using a sliding window of 120 codon sequences and found that the enzyme active center regions indeed fell within sequences under positive selection.

Considering the multiple influences of genetic code structure and amino acid physicochemical properties on protein evolution, we calculated the ratio of amino acid sites differing by more than 1 Å to those below this threshold (named the structural conservation index, SCI) to characterize protein structural conservatism, specifically, align pairwise protein structures using US-align with a threshold set to -d 1 Å. The number of amino acids marked by ":" (denote residue pairs with d < 1 Angstrom) was extracted. (denote other aligned residues), and gaps, labeled NPR, OAR, and NMR. The SCI was calculated as NPR/AR - OAR/AR - 2*NMR/AR, where AR represents the total number of sites. The range of the SCI is from -2 to 1, and the closer the value is to 1, the more similar the two protein structures are.

**Prediction of Protein Subcellular Localization**

The cellular localization of proteins from the JK6 strain and those reconstructed from ancestors was predicted using a combination of protein localization prediction software, including PSORTb v3.0[31], Cell-PLoc 2.0[32], CELLO v.2.5[33], and LocTree3[34], in conjunction with SignalP 6.0[35]. Additionally, the distinction was aided by comparing the ratio of supernatant to total protein content in the cell culture fluid.

**Evaluation of gene–gene covariation, pairwise identity and evolutionary rate**

We extracted species from the collected genomic dataset that had homologs of all four target proteins. Then, we constructed a species tree by merging the common genes of all these species that were aligned using MAFFT[36]. Subsequently, constrained gene trees were built using the species tree as a reference constraint (with iqtree2[37], specifying the species tree using the -keep-ident -te parameter). To calculate gene–gene covariation, we used phykit[38] with the command "cover -r" to reference the species tree. The evolutionary rate was determined using "evolutionary_rate", and pairwise identity was calculated using "pairwise_identity".

**The expression and purification of recombinant proteins**

Protein Expression and Purification. The genes corresponding to agarase were amplified by PCR from the genomic DNA of the JK6 strain. The sequences of primers used for cloning are listed in Table S1. Each amplified DNA fragment was inserted into the EcoRI-XhoI site of pPEI-His-Sumo. *Escherichia coli* strain BL21(DE3) cells (Vazyme) carrying each expression plasmid were cultured in 2 L of LB media supplemented with 50 μg/mL carbenicillin at 37°C and induced with 1 mM isopropyl β-D-1-thiogalactopyranoside (A100487, Sangon Biotech) at 16°C for 16 h when the OD600 reached 0.8. The cells were subsequently harvested by centrifugation at 4000 × g for 10 min at 4°C. The cell pellet was resuspended in 80 mL of lysis buffer (20 mM Tris-HCl pH 8.0, 500 mM NaCl, 5 mM imidazole) supplemented with 1 mM phenylmethanesulfonyl fluoride (PMSF, A100754, Sangon Biotech). Then, the cells were disrupted by a high-pressure homogenizer (ATS Engineering Limited, AH1500) for 3 min at 800 bar, and the extracted proteins were separated from the cell debris by centrifugation at 12000 × g for 1 h at 4°C. His-tagged proteins were purified using a HisTrap HP column (5 mL, Cytiva) with a Fast Protein Purification System (AKTA Pure 25) and eluted with elution buffer (20 mM HEPES pH 7.5, 300 mM imidazole, 100 mM KCl). The proteins were further enriched using a HiTrap Q HP Column (5 mL, Cytiva) and separated by gradient elution with buffer A (20 mM HEPES pH 7.5, 50 mM KCl, 1 mM DTT) and buffer B (20 mM HEPES pH 7.5, 1 M KCl, 1 mM DTT). The peak fractions containing the agarase protein were collected at approximately 10 mg/mL for further experiments.

The gene sequences of the *Vibrio astriarenae* strain (HN897) strain were synthesized by GenScript, and the strains were named *Vsp6*, *Vsp11*, *Vsp12* and *Vsp18* based on their homology with the four agarase proteins in JK6. Following construction through standard methods for sequence replacement and point mutation in plasmids, the procedures were executed consistently with the previously described conventional protein expression process. The fragment "FADKDKRSGKEVASEV" in PL3512 was replaced by "YDHSKVAARSDDDVTPEDSKEKLEISQEAFDSAYVANQT", which was derived from the PL3506 fragment. The fragment "STSGADRLPVKQIGSG" in PL3511 was replaced by "IFGDHARISTGND", which was derived from PL3512. The following active parts, "HSKVAARSDDDVTPEDSKEKLEISQEAFDSAY" in PL3506 and "LSEDLSLRPSQISQDSYVDNQ" in PL3518, were deleted from their sequences. The modified proteins were marked as PL3506-M, PL3511-M, PL3512-M and PL3518-M. The point mutant of PL3506 was named 6 M, the glycoside hydrolase domain of PL3506 was named 6 g, and the point mutation in the glycoside hydrolase domain of PL3506 was named 6gM. The amino acid sequences of each protein and the modified versions are shown below.

**Agonase activity assay**

The activity of the agarases was measured using the 3,5-dinitrosalicylic acid (DNS) method[39]. Briefly, 10 µL of each crude enzyme sample was added to 490 µL of PBS (pH 7.4) containing 0.1% agarose powder and incubated at 37°C for 1 hour. After centrifugation at 12,000 rpm, 50 µL of the supernatant was added to a 96-well PCR plate containing 100 µL of DNS reagent. As controls, equal volumes of D-galactose solutions with gradient concentrations were added to other wells. The PCR plate was sealed with adhesive film and centrifuged to the bottom. Then, the plate was heated at 99°C for 10 minutes in a PCR machine and immediately cooled on ice to stop the reaction. After cooling, 100 µL of each well solution was transferred to an enzyme-linked immunosorbent assay (ELISA) plate. The absorbance was measured at 540 nm using an ELISA reader (Thermo Varioskan LUX). The concentration of reducing sugars produced by each enzyme was calculated based on a standard curve.

**Analysis of the enzymatic hydrolysis products using thin-layer chromatography (TLC)**

The products of agarose hydrolysis by various agarases were separated using thin-layer chromatography (TLC)[39-41]. After expression and purification, the crude enzyme samples (20 µL) were digested overnight at room temperature in 500 µL of PBS (pH 7.4) containing an excess of agar powder. Following centrifugation at 12,000 rpm, the supernatant was collected, and aliquots of 5 µL were spotted onto silica gel 60 plates (Merck Millipore) and dried prior to chromatography. Two different mobile phases and staining solutions were used for chromatographic separation and visualization, respectively. The first mobile phase consisted of 1-butanol/acetic acid/distilled water (2:1:1, v/v/v)[42], and the gels were visualized with a 3 mL/40 mL sulfuric acid/ethanol solution, followed by heating in a microwave oven for 30 seconds. The second mobile phase was composed of 1-butanol/acetic acid/distilled water (6:3:2, v/v/v), with visualization achieved using a 3 mL/40 mL sulfuric acid/ethanol solution containing 0.2% 1,3-dihydroxynaphthalene[43], followed by a similar heating step in a microwave oven for 30 seconds. As standards, NA2, A3, NA4, and agarose oligosaccharides were purchased from Qingdao Bozhi Huli Biotechnology Co., Ltd., and Gal was purchased from Sangon Biotech Co., Ltd.

**Mass Spectrometry Techniques for Polysaccharide Analysis**

To identify the molecular weights of sugars from different bands obtained via thin layer chromatography (TLC), the major bands were scraped, combined, and dissolved in deionized water (ddH2O) before centrifugation to collect the supernatant. The hydrolysis products from different proteins were filtered directly through a 5 kDa ultrafiltration tube to measure the filtrate. All samples were sent to The Mass Spectrometry & Metabolomics Core Facility at Westlake University for analysis using quadrupole time-of-flight mass spectrometry (Q/TOF-MS). The specific process involved the following steps: The MS and Metabolomics Core Facility at Westlake University was provided with the combined major bands obtained from TLC, dissolved in ddH2O and then centrifuged to obtain the supernatant. The sample was subjected to Q/TOF-MS mass spectrometry detection. The specific process was as follows: chromatographic elution was performed on an Agilent 1290 infinity system (Agilent Technologies, USA) equipped with a sample manager coupled to a mass spectrometer with an electrospray ion source in both positive and negative ion modes. The separation was carried out on a BEH amide column (100 mm × 2.1 mm, 1.7 μm) at 40°C. Mobile phase A was ultrapure water containing 0.3% ammonia and 15 mM ammonium acetate, and mobile phase B was acetonitrile/water (9:1) containing 0.3% ammonia and 15 mM ammonium acetate. The flow rate was 0.3 mL/min, and the gradient of mobile phase A was held at 5% for 1 min, 5% to 50% for 8 min, 50% for 3 min, 50% to 5% for 0.5 min, and finally held at 5% for 6.5 min. The sample volume injected was 5 μL.

An Agilent 6545 Q/TOF-MS system in both the ESI+ and ESI- modes was used for detection. The parameters used were as follows: capillary voltage, 3500 V; nozzle voltage, 120 V; nebulizer gas, 35 psi; drying gas flow rate, 8 L/min; and gas temperature, 350°C. A full scan was run with a mass range from m/z 50 to 1200 using high-resolution mode (Extended Dynamic Range 2 GHz). Additionally, to produce accurate mass correction, an automated calibration delivery system was used to calibrate the MS and MS/MS results automatically by prepared calibration solutions before and during the analysis. The data were processed using a qualitative workflow.

**Proteomic analysis of the JK6 strain.**

In this study, protein samples were prepared for mass spectrometry analysis in two groups. For the first group, bacterial cells were cultured in L1A media for 24 hours, collected by centrifugation, and washed twice before being inoculated into equal volumes of L1A, L1B, or L1 media. After another 24 hours of cultivation, the cells were lysed under high pressure, and the supernatant containing proteins was collected for analysis. The second group followed a similar procedure, starting with cultivation in L1B media. However, after the second 24-hour cultivation in L1A, L1B, and L1 media, both the cells and the culture supernatant were collected. The culture supernatant was concentrated using a 10-kD concentration tube, and the cells were lysed under high pressure to collect and concentrate the proteins in the supernatant. This approach enabled the comparison of protein expression profiles under different culture conditions, providing valuable insights into the impact of these conditions on cellular protein production and secretion. Then, mass spectrometry detection was performed.

SDS–PAGE was used to separate the proteins, which were then stained with Coomassie Blue G-250. The gel bands of interest were cut into pieces. The sample was digested with trypsin prior to reduction and alkylation in 50 mM ammonium bicarbonate at 37°C overnight. The digested products were extracted twice with 1% formic acid in 50% acetonitrile aqueous solution and then dried using a SpeedVac.

The extracted protein was quantified using BCA assays (Thermo Scientific). After reduction, alkylation, and precipitation with acetone, 50 µg of protein was digested using trypsin at 37°C overnight. The digested products were desalted using solid-phase extraction cartridges and then subjected to LC–MS analysis.

For LC–MS/MS analysis, the peptides were separated by 65 min of gradient elution at a flow rate of 0.300 µL/min with a Thermo EASY-nLC1200 integrated nano-HPLC system, which is directly interfaced with a Thermo Q Exactive HF-X mass spectrometer. The analytical column was a homemade fused silica capillary column (75 µm ID, 150 mm length; Upchurch, Oak Harbor, WA) packed with C-18 resin (300 A, 3 µm, Varian, Lexington, MA). Mobile phase A consisted of 0.1% formic acid, and mobile phase B consisted of 100% acetonitrile and 0.1% formic acid. The mass spectrometer was operated in the data-dependent acquisition mode using the Xcalibur 4.1 software, and there was a single full-scan mass spectrum in the Orbitrap (400-1800 m/z, 60,000 resolution) followed by 20 data-dependent MS/MS scans at 30% normalized collision energy. Each mass spectrum was analyzed using the Thermo Xcalibur Qual Browser and Proteome Discovery for database searching.

**Reverse transcription and quantitative real-time PCR (RT–qPCR)**

The sample processing involved culturing bacterial cells in L1B medium for 24 hours, followed by transfer to L1 medium for an additional 36 hours. After centrifugation and resuspension, the samples were collected at 0 hours and then transferred to various modified L1 media—agar-L1 media containing 5% agar, GAL-L1 media supplemented with 0.1 g/L galactose, NA2-L1 media supplemented with 0.1 g/L NA2, NA4-L1 media supplemented with 0.1 g/L NA4, S-L1 media supplemented with 0.1 g/L agar oligosaccharides, and S10-L1 media supplemented with 1 g/L agar oligosaccharides—with three bottles per cultivation condition for each replicate. The agar oligosaccharides, as shown by thin layer chromatography (TLC), had a chain length greater than that of NA8 but were uniformly soluble in the culture medium. Timing began upon transfer to the corresponding media, with samples collected at 5 minutes, 30 minutes, 1 hour, 5 hours, and 20 hours posttransfer, followed by rapid freezing in liquid nitrogen.

Based on previous RNA sequencing results, the internal reference genes PL0252 and PL1865 were selected for qPCR analysis. A single pair of amplification primers was designed for each target gene, with three pairs of primers designed for certain genes of special interest. According to the manufacturer's instructions, RNA from the samples was extracted using the FastPure Cell/Tissue Total RNA Isolation Kit V2 produced by Nanjing Vazyme Biotech Co., Ltd. Subsequently, HiScript III RT SuperMix for qPCR (+gDNA wiper) R323 (Vazyme) was used for reverse transcription. ChamQ Universal SYBR qPCR Master Mix Q711 (Vazyme) was used for each qPCR.

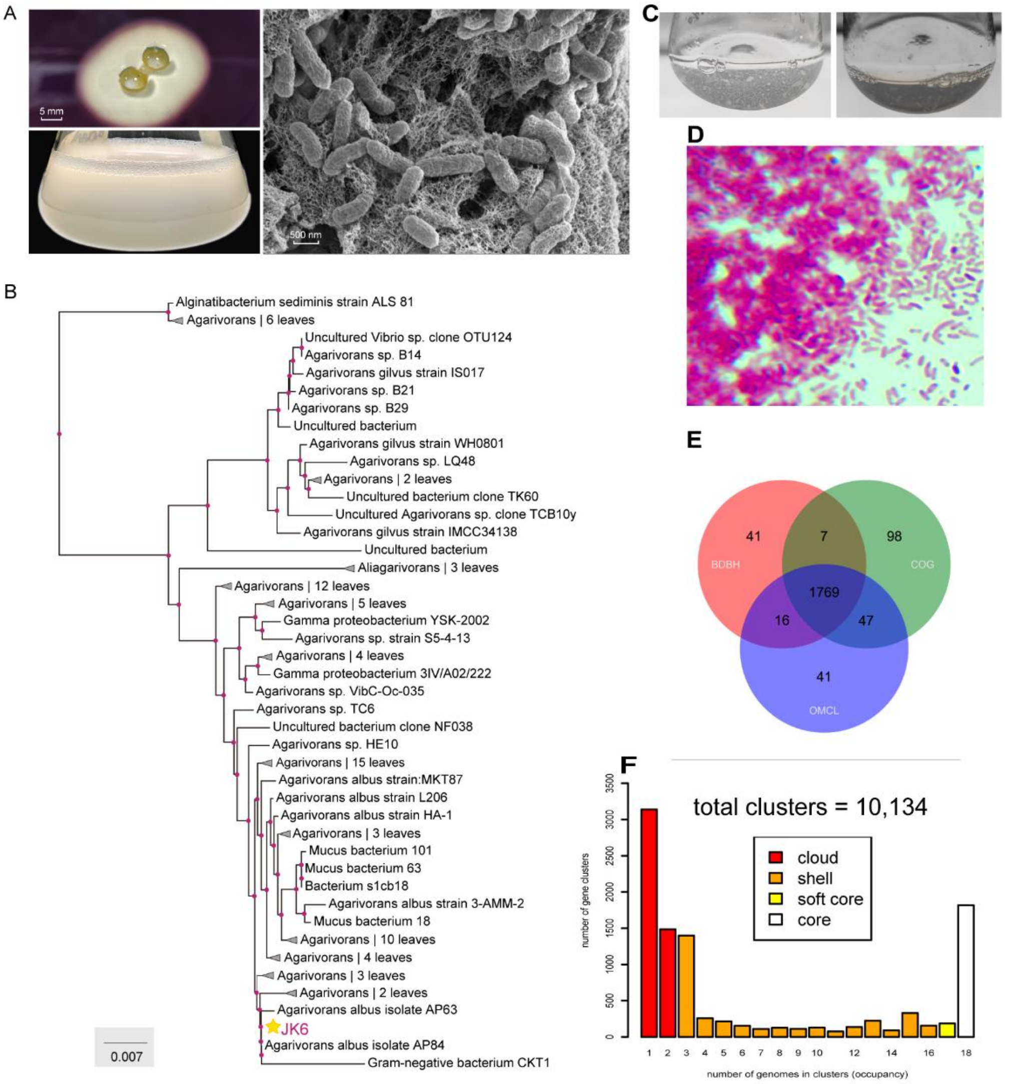


**Fig. S1. Physiological Culturing Characteristics and Pangenome Analysis of the JK6 Strain**
**(A)** The strain formed hydrolysis circles on L1 medium plates with transparent staining and grew rapidly and in large quantities in L1B medium, reaching concentrations on the order of $1\times10^{9}$. Scanning electron microscopy revealed that the bacteria were short rod-shaped, had a wrinkled surface, and closely adhered to the agar surface. **(B)** Amplification and sequencing of the 16S rRNA sequence of the JK6 strain followed by tree construction using NCBI BLAST identified it as a member of the genus *Agarivorans*, and together with the growth phenotype, confirmed it to be *Agarivorans albus*. **(C)** After culturing overnight in L1A liquid medium supplemented with 0.2% agar, the bacteria clumped into white patches, and the solution became clear and transparent after one week. **(D)** Gram-stained bacteria were negative. **(E, F)** Pangenomic comparison detailing the genes shared among 17 closely related strains within the same genus and the JK6 strain.

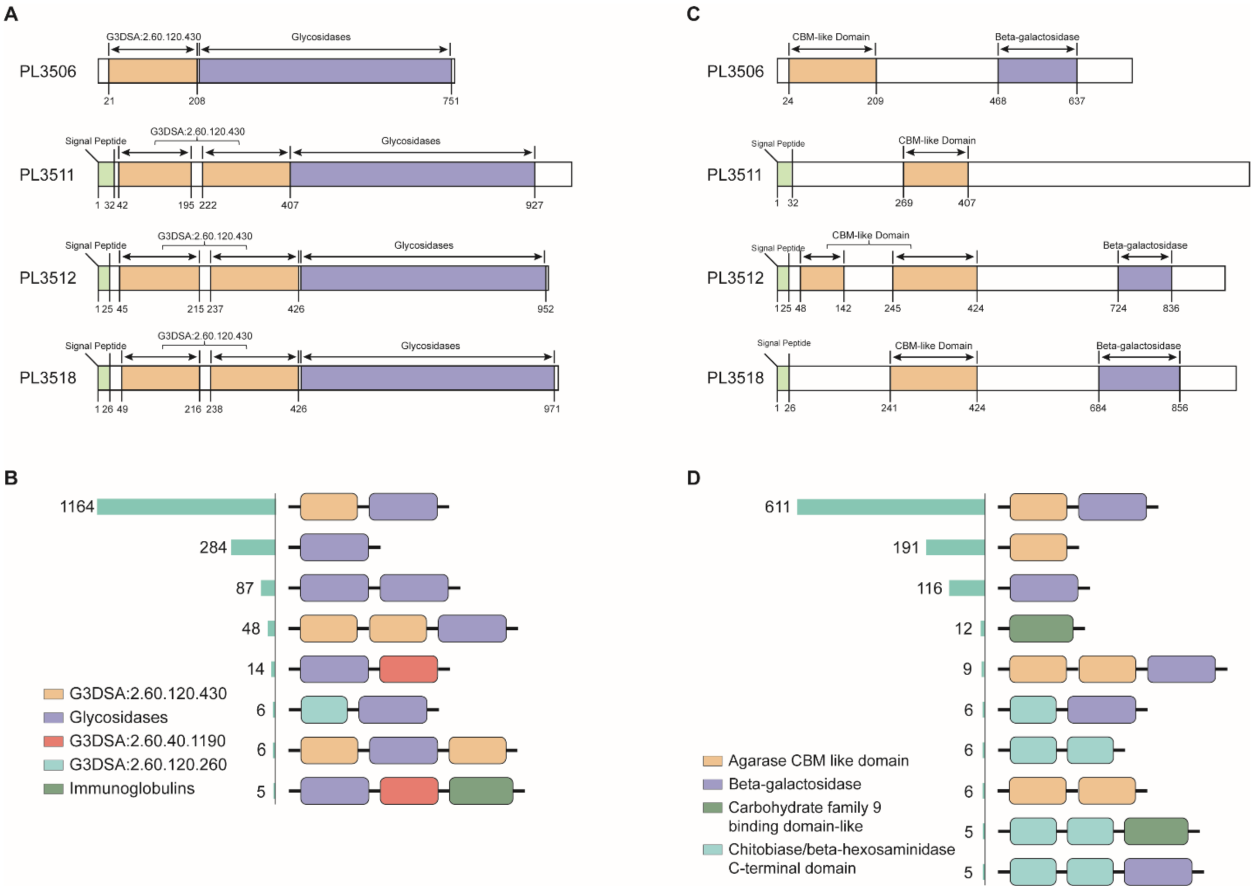


**Fig. S2. Gene3D and Pfam annotations reveal the structural domain scenarios for the four proteins and genes of the GH50 family.** **(A)** Gene3D annotation of the structural domains of four genes. **(B)** Gene3D annotation of the structural domains of the GH50 family. **(C)** Pfam annotation of the structural domains of the four genes. **(D)** Pfam annotation of the structural domains of the GH50 family genes. In terms of gene structure, despite differences in annotations between Gene3D and Pfam, the AlphaFold-predicted structures suggested that the predictions made by Gene3D were more accurate. According to the annotation statistics for GH50 family proteins, the combination of dual CBM-like domains (G3DSA:2.60.120.430) with a glycoside hydrolase domain is relatively rare in the GH50 family.

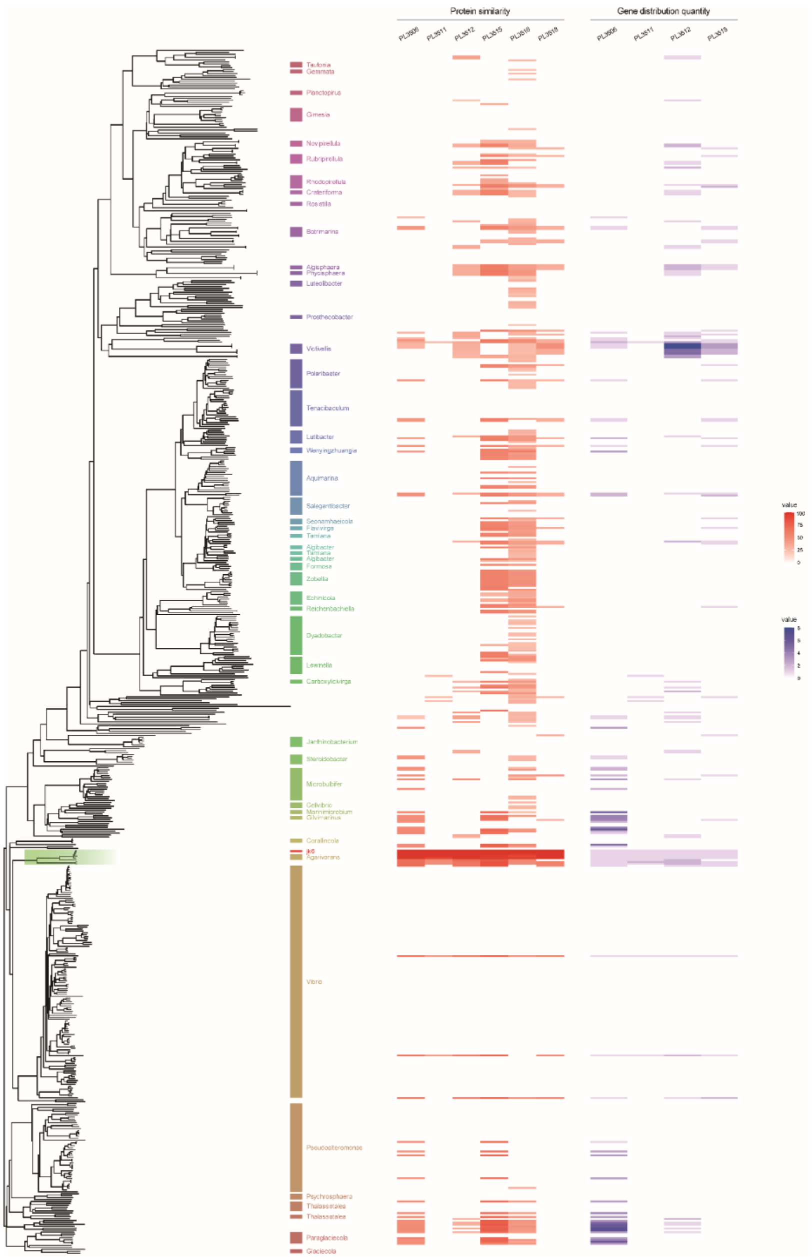


**Fig. S3. The number of homologous genes and the highest homology of four proteins in 629 bacterial strains.** On the left is the phylogenetic tree, with text annotations indicating the topological structure showing three or more consecutive genus names. In the middle are the highest protein similarities for the four proteins and PL3515/PL3516 within the gene clusters of the four proteins. On the right is the number of occurrences of the four proteins in each species. The homologs of these four proteins vary in quantity across different clades according to evolutionary relationships. The homologous genes of proteins PL3506 and PL3512 are primarily distributed in different clades and often do not coexist in the same species. Based on the inferred phylogenetic topology of the four genes, we observed that the earliest diverged PL3511 had the least number of homologs across species, while the number of homologs in transitional PL3518 was slightly greater but still significantly less than that in PL3506 and PL3512.

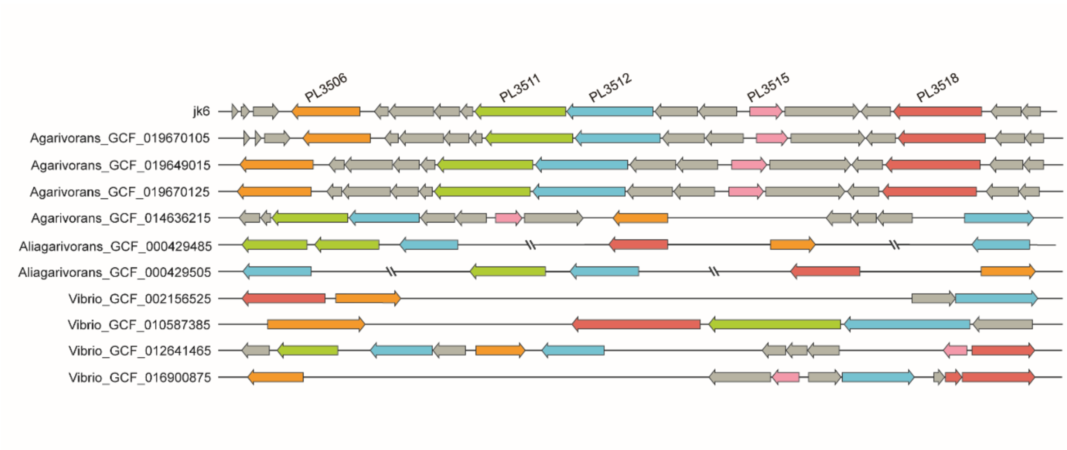


**Fig. S4. The gene cluster structures in several representative species containing the four proteins.** The four proteins and PL3515 are marked with different colors, and double slashes indicate discontinuities in the genome. The gene structures were largely consistent within the genus *Agarivorans*, and the genes PL3511 and PL3512 still retained a tandem duplication pattern adjacent to each other, although structural changes occurred in one strain. In strains from other genera, significant structural variations are evident.

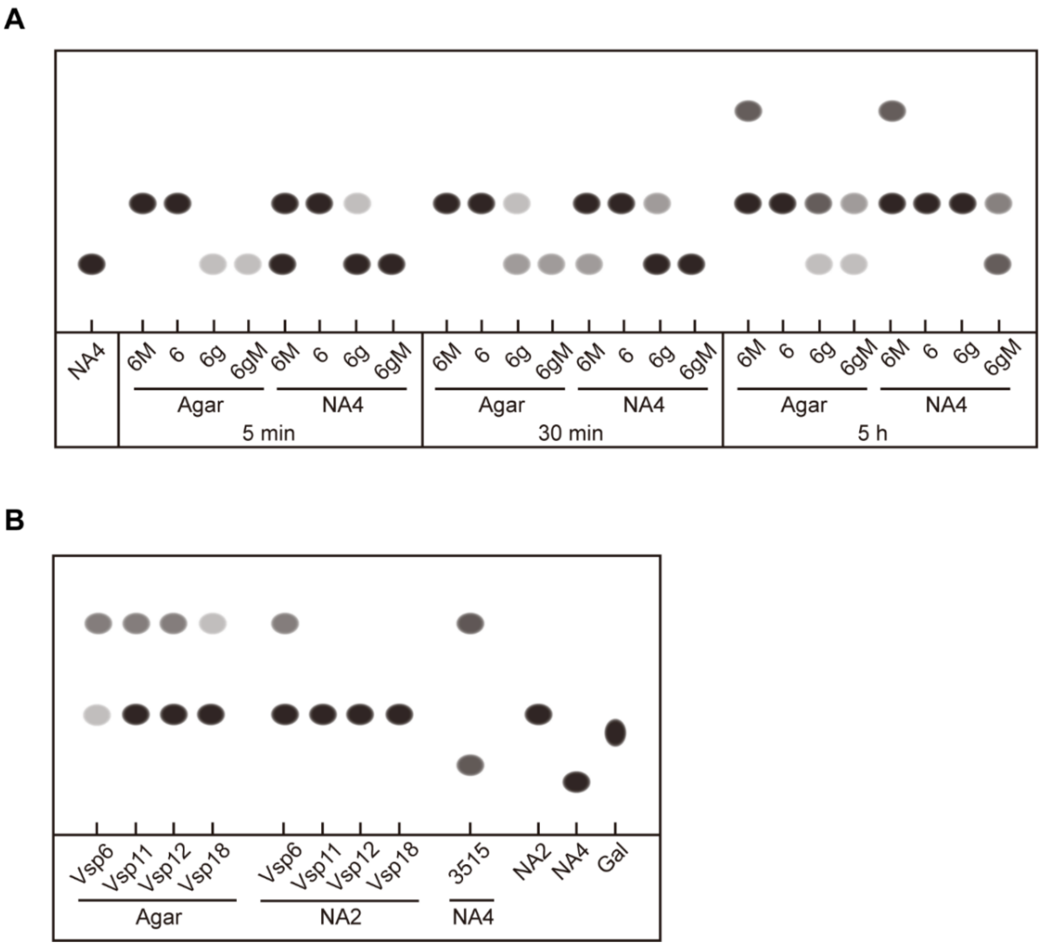


**Fig. S5. Hydrolytic activity of mutants of the PL3506 protein and the four homologous proteins in the *Vibrio astriarenae* strain. (A)** 6 M represents a mutant protein obtained by point mutation of the complete PL3506 protein, 6 represents the complete PL3506 protein, 6 g represents the protein expressed by the glycoside hydrolase domain of PL3506, and 6gM represents the protein of the same site point mutation in the glycoside hydrolase domain of PL3506. 6 M can hydrolyze both agar and NA4 into monosaccharide products over time, but the hydrolysis rate of NA4 is significantly slower than that of the original protein. 6 g has a slower rate of hydrolyzing NA4 and produces NA4 bands during the hydrolysis of agar, which were not present in the original protein. The hydrolytic activity of 6gM is similar to that of 6 g, but the hydrolysis rate of NA4 is significantly slower than that of 6 g. Within the observed time, NA4 in the substrate is not completely consumed, leaving it uncertain whether it also produces monosaccharide hydrolysis products similar to 6 M. **(B)** Vsp6 to Vsp18 represent the homologous proteins in *Vibrio astriarenae* (HN897) corresponding to PL3506-PL3518. Among them, all four proteins in HN897 produced weak monosaccharide bands, but in comparison to the hydrolysis of NA2, except for Vsp3506, which can hydrolyze NA2, the other three proteins did not have this activity, indicating that their monosaccharide production is due to other activities. The grayscale spots in the image represent hydrolysis products, with the depth of grayscale indicating the concentration of the products.

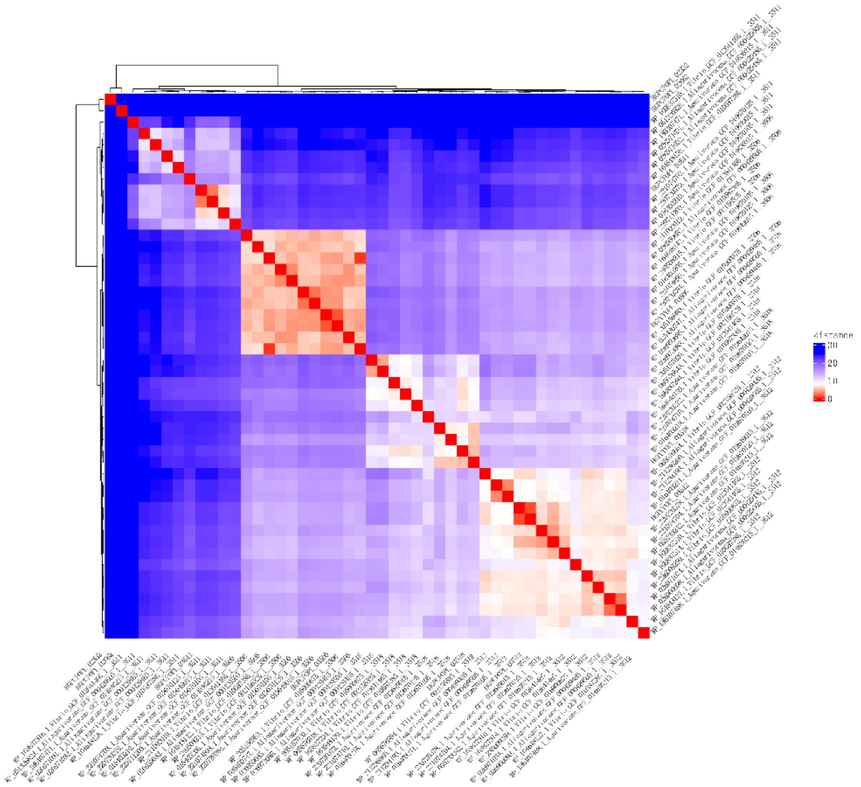


**Fig. S6. Glycoside hydrolase domain structural similarity clustering heatmap.** A heatmap was constructed using the Vd values calculated from the eigenvalues obtained through the alignment of the glycoside hydrolase domains of the compared proteins using USalign. The heatmap shows that the four proteins are distinctly divided into four units.

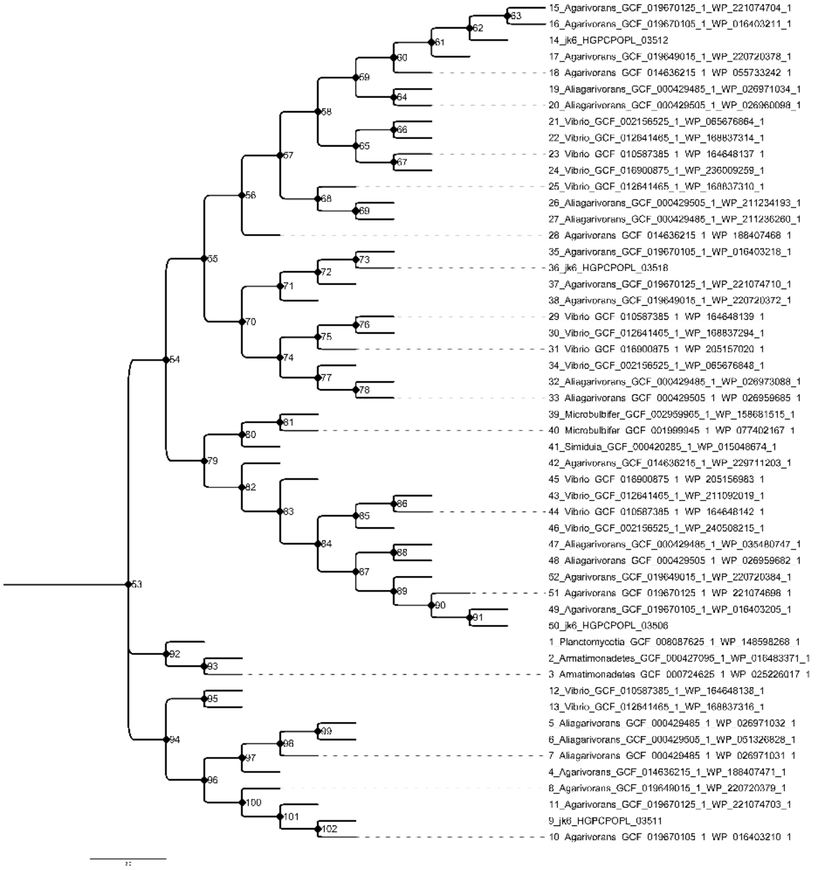


**Fig. S7. PAML-reconstructed ancestral tree of the four proteins.** The ancestors related to PL3511, PL3512, and PL3518 essentially maintained the same protein localization as their corresponding contemporary proteins. However, the ancestral proteins of PL3506 showed different localization patterns. The numbers marked at each node represent the ancestral protein numbers, consistent with the numbering in the ancestral protein localization table (Data S6).

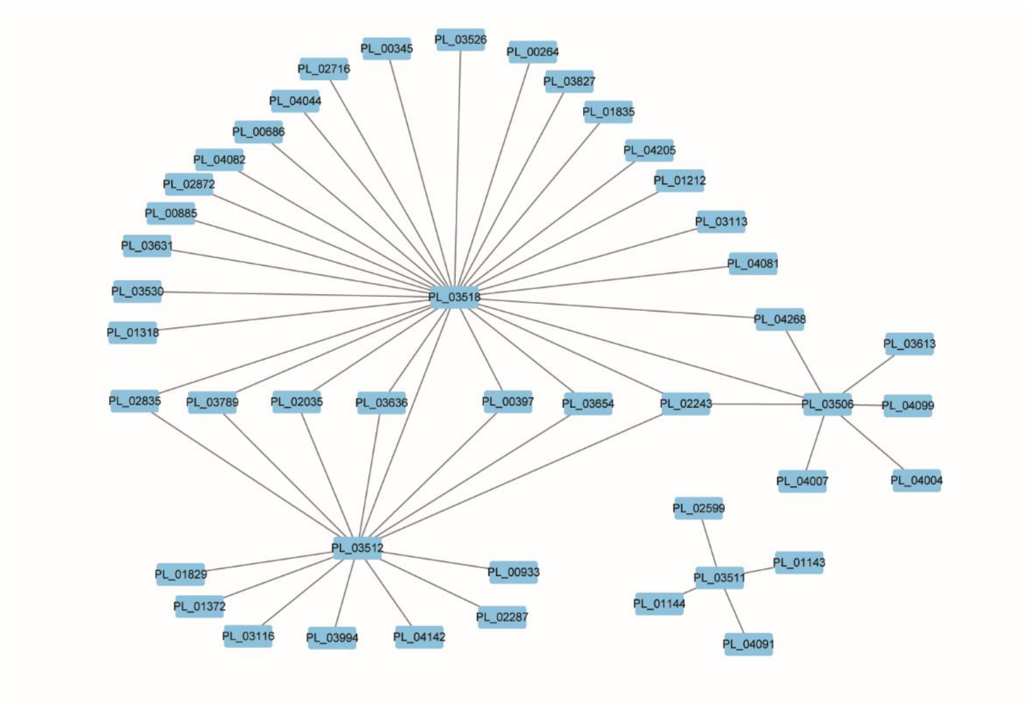


**Fig. S8. Coevolutionary history of the four proteins and the shared proteins among the collected species.** Covarying evolutionary rates between pairs of proteins, calculated using PhyKit, with a threshold of 8.0, were used to filter protein pairs related to the four proteins. Within the set threshold range, we found that PL3511 did not exhibit a significant correlation with the other three agarase genes (PL3506, PL3512, and PL3518). However, PL3511 shows a covariation relationship with enzymes related to sugar metabolism, such as carbonic anhydrase and glycosyltransferase. Among PL3506, PL3512, and PL3518, we discovered that they are related to a commonly associated gene, the glycoside/cation symporter (PL2243). Additionally, the strongest association of PL3506 was with the type IV pilin protein (PL3613), which is related to host adhesion and is associated with the greatest number of proteins related to phosphate metabolism processes. For PL3512, the protein with the highest association was the periplasmic transport-related protein PL1829 (Tol-Pal system protein TolQ), which matches its cell localization and is associated with the greatest number of proteins related to serine hydrolase activity and peptidase activity. Moreover, PL3518 had the strongest association with PL3512, followed by the protein PL2243, which is related to all three enzymes; the strongest independent association was with probable peptidoglycan glycosyltransferase (PL3631), which is associated with the greatest number of proteins related to catalytic activity and hexosyltransferase activity. A network diagram was constructed using Cytoscape to visualize the associations.

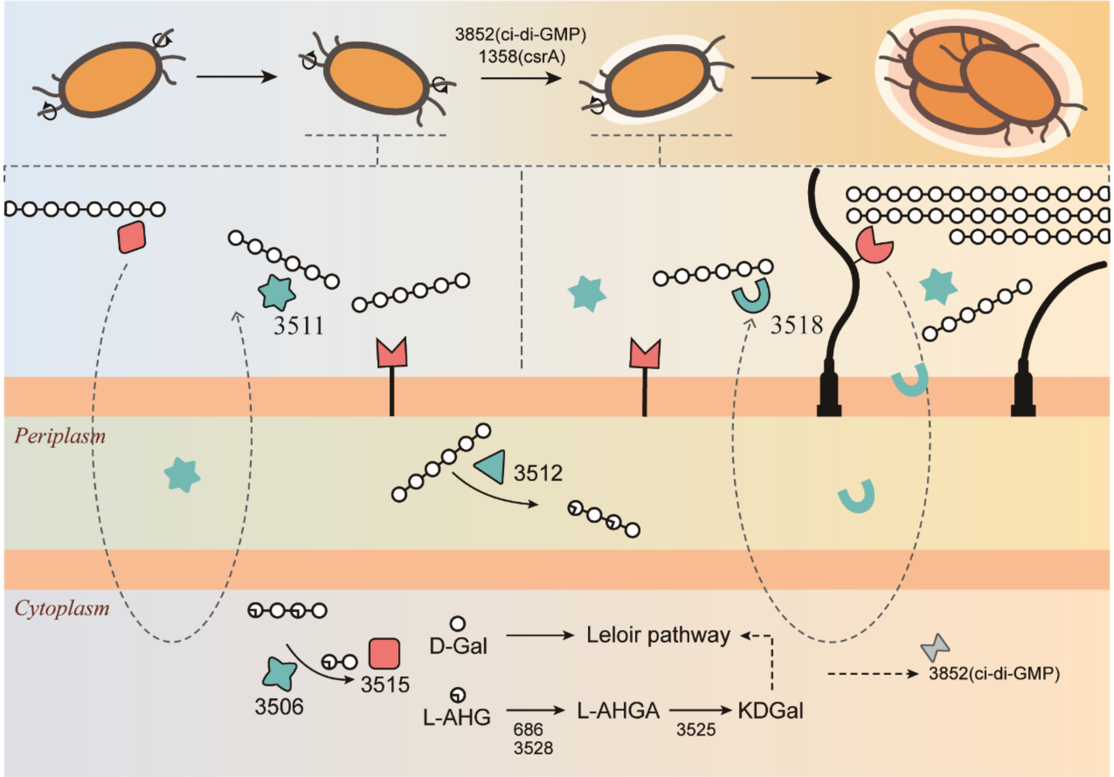


**Fig. S9. Regulatory model of the perception and response of the JK6 strain to agar induction.** Macroscopically, upon sensing the presence of agar and its abundance in the environment, the bacteria transition from individual swimming states to multicellular aggregation, forming biofilms that adhere to the surface of agar bodies. At the molecular level, in the free-swimming state, certain agarases are secreted extracellularly to explore and sense the presence of agar in the environment. When encountering low levels of agar, certain agarases are predominantly expressed at specific ratios for the degradation and absorption of agar. Conversely, in the presence of high concentrations of agar, other agarases take precedence or are expressed in different ratios to regulate the absorption and utilization of agar. Intracellular PL3506 hydrolyzes it into disaccharides, which are then hydrolyzed into monosaccharides by PL3515, which subsequently enter metabolic pathways such as AHG metabolism and the Leloir pathway. The process of agar absorption and utilization is accompanied by the induction of signals such as ci-di-GMP, leading to the transition of bacteria from a free-swimming state to an attached state.

**Table S1.**

List of primers used in the experiment.

| ID # | Assay | Description | Sequences (5'-3') |
|---|---|---|---|
| PL2332.FOR | Recombinant protein expression in E.coli | Forward primer for PL2332 cloning into pPEI-His-Sumo | GCACCCACACGCGCGAATTCATGACTATCAAGGCAAAGAAAATGGC |
| PL2332.REV | Recombinant protein expression in E.coli | Reverse primer for PL2332 cloning into pPEI-His-Sumo | TGGTGGTGGTGGTGCTCGAGTTAGCGTTGTGGCGGGGC |
| PL3529.FOR | Recombinant protein expression in E.coli | Forward primer for PL3529 cloning into pPEI-His-Sumo | GCACCCACACGCGCGAATTCATGAAAGGATTCACTAAGCACCC |
| PL3529.REV | Recombinant protein expression in E.coli | Reverse primer for PL3529 cloning into pPEI-His-Sumo | TGGTGGTGGTGGTGCTCGAGCTACTGGAATTTAAAACGTTGGTTGG |
| PL3506.FOR | Recombinant protein expression in E.coli | Forward primer for PL3506 cloning into pPEI-His-Sumo | GCACCCACACGCGCGAATTCATGGAAACGCGCTTAAGGGC |
| PL3506.REV | Recombinant protein expression in E.coli | Reverse primer for PL3506 cloning into pPEI-His-Sumo | TGGTGGTGGTGCTCGAGCTACTTTTTCGCCTTATTAAAACGG |
| PL0687.FOR | Recombinant protein expression in E.coli | Forward primer for PL0687 cloning into pPEI-His-Sumo | GCACCCACACGCGCGAATTCATGCTGTTTAGAAACACTCTATTACTTG |
| PL0687.REV | Recombinant protein expression in E.coli | Reverse primer for PL0687 cloning into pPEI-His-Sumo | TGGTGGTGGTGGTGCTCGAGTTAATTTACGAATTTAAAGCTTTGA |
| PL3512.FOR | Recombinant protein expression in E.coli | Forward primer for PL3512 cloning into pPEI-His-Sumo | GCACCCACACGCGCGAATTCATGGTAGAAGTTATGAAATTTACAAAAAAT |
| PL3512.REV | Recombinant protein expression in E.coli | Reverse primer for PL3512 cloning into pPEI-His-Sumo | TGGTGGTGGTGGTGCTCGAGTTATTTTTTGTAACGAAGCATGTAAACG |
| PL2878.FOR | Recombinant protein expression in E.coli | Forward primer for PL2878 cloning into pPEI-His-Sumo | GCACCCACACGCGCGAATTCATGCAGTTCATGTTAAAGCGCC |
| PL2878.REV | Recombinant protein expression in E.coli | Reverse primer for PL2878 cloning into pPEI-His-Sumo | TGGTGGTGGTGGTGCTCGAGCTATTGGCAAGTATAACCTGATACAAC |

| | | | |
|---|---|---|---|
| PL3511.FOR | Recombinant protein expression in E.coli | Forward primer for PL3511 cloning into pPEI-His-Sumo | GCACCCACACGCGCGAATTCATGAAGATTAAATTTTTATCTGCAGC |
| PL3511.REV | Recombinant protein expression in E.coli | Reverse primer for PL3511 cloning into pPEI-His-Sumo | TGGTGGTGGTGGTGCTCGAGTTACACTTTACGACGTCTTAGTAAAA |
| PL2076.FOR | Recombinant protein expression in E.coli | Forward primer for PL2076 cloning into pPEI-His-Sumo | CACCCACACGCGCGAATTCATGGAATATATGATGATTCGTAAAAATAATC |
| PL2076.REV | Recombinant protein expression in E.coli | Reverse primer for PL2076 cloning into pPEI-His-Sumo | TGGTGGTGGTGGTGCTCGAGTTATTCGTCAACGAGTTTCCAAG |
| PL3518.FOR | Recombinant protein expression in E.coli | Forward primer for PL3518 cloning into pPEI-His-Sumo | GCACCCACACGCGCGAATTCTTGCTTAGTGGTTGTCCAAATATTG |
| PL3518.REV | Recombinant protein expression in E.coli | Reverse primer for PL3518 cloning into pPEI-His-Sumo | TGGTGGTGGTGGTGCTCGAGCTACTTCACCAGCTCTGGGTAAC |
| PL3515.FOR | Recombinant protein expression in E.coli | Forward primer for PL3515 cloning into pPEI-His-Sumo | GCACCCACACGCGCGAATTCATGTCAGGGACAGGTAGCAAG |
| PL3515.REV | Recombinant protein expression in E.coli | Reverse primer for PL3515 cloning into pPEI-His-Sumo | TGGTGGTGGTGGTGGTGCTCGAGTTAATTTGAATTTTGATAAGTTGAAGC |
| PL3512M.R1 | Recombinant protein expression in E.coli | primer for modified protein PL3512-M expression construction | GTGGTCGTAGTCGTGGCCAGTAATGGTTAC |
| PL3512M(+06).F | Recombinant protein expression in E.coli | primer for modified protein PL3512-M expression construction | CTGGCCACGACTACGACCACTCAAAAGTGGC |
| PL3512M(+06).R | Recombinant protein expression in E.coli | primer for modified protein PL3512-M expression construction | TTGAACGGCGAGTTTGGTTAGCAACATAGG |
| PL3512M.F2 | Recombinant protein expression in E.coli | primer for modified protein PL3512-M expression construction | TAACCAAACTCGCCGTTCAATGTTTACTTG |
| PL3506M.R1 | Recombinant protein expression in E.coli | primer for modified protein PL3506-M expression construction | GGTTAGCAACGTCGTAATCAATACCGGTAA |

| | | | |
|---|---|---|---|
| PL3506M.F2 | Recombinant protein expression in E.coli | primer for modified protein PL3506-M expression construction | TGATTACGACGTTGCTAACCAAACTCGCCG |
| PL3518M.R1 | Recombinant protein expression in E.coli | primer for modified protein PL3518-M expression construction | AACGAGCATCCACCAAATCGGTAAAGCCAA |
| PL3518M.F2 | Recombinant protein expression in E.coli | primer for modified protein PL3518-M expression construction | CGATTTGGTGGATGCTCGTTCAGTGGCTTC |
| PL3511M.R1 | Recombinant protein expression in E.coli | primer for modified protein PL3511-M expression construction | AGTGCTAATGCGTGCGTGGTCTCCAAAAATCCAACCATTTGCCACGTAAG |
| PL3511M.F2 | Recombinant protein expression in E.coli | primer for modified protein PL3511-M expression construction | ACCACGCACGCATTAGCACTGGTAACGACTACTGGGGACCACTTCCTGA |
| PL3506g.FOR | Recombinant protein expression in E.coli | Forward primer for PL3506 glycoside hydrolase domain cloning into pPEI-His-Sumo | GCACCCACACGCGCGAATTCGGTTGTTTAGATAAGTTTGG |
| PL3506g.REV | Recombinant protein expression in E.coli | Reverse primer for PL3506 glycoside hydrolase domain cloning into pPEI-His-Sumo | GGTGGTGGTGGTGCTCGAGTTAACGGCGCTGATACATGCC |
| PL3506gM.F | Recombinant protein expression in E.coli | Primer for point mutant of PL3506 glycoside hydrolase domain | TTGATAATGGGAAAAGCTGGGGCCGAATGG |
| PL3506gM.R | Recombinant protein expression in E.coli | Primer for point mutant of PL3506 glycoside hydrolase domain | CAGCTTTTCCCATTATCAATGAAAATACCC |
| 0252qF | qPCR | qPCR primer of PL0252 | TGGTCACTCAACAAGCCTCT |
| 0252qR | qPCR | qPCR primer of PL0252 | AGCAAAGGCATCAAGGCAAA |
| 1865qF | qPCR | qPCR primer of PL1865 | TTGTTGGCGGAATACGTCAC |
| 1865qR | qPCR | qPCR primer of PL1865 | ACTGCCAGTACAGCAGTGAT |
| 3506qF | qPCR | qPCR primer of PL3506 | CGATTACTGGGCGTTCGTTT |

| | | | |
|---|---|---|---|
| 3506qR | qPCR | qPCR primer of PL3506 | TACATGCCGCCATTTAACCG |
| 3506qF2 | qPCR | qPCR primer of PL3506 | CAGATCCTTTCGACCCGGTA |
| 3506qR2 | qPCR | qPCR primer of PL3506 | TGGTGAACCTTGCATTTCGG |
| 3511qF | qPCR | qPCR primer of PL3511 | TATGCAAACGGTGACGTTCC |
| 3511qR | qPCR | qPCR primer of PL3511 | TAGCCGCTGCCAATTTGTTT |
| 3511qF2 | qPCR | qPCR primer of PL3511 | GATGGCGAAACAGTCACTCC |
| 3511qR2 | qPCR | qPCR primer of PL3511 | CCCACCATCAGGTCGTAAGT |
| 3511qF3 | qPCR | qPCR primer of PL3511 | ATTATGCGTCTTGGGATGCG |
| 3511qR3 | qPCR | qPCR primer of PL3511 | AGCAAATTGCCAAGCAAGGA |
| 3512qF | qPCR | qPCR primer of PL3512 | ACCCACAAGGCAACTTGTTC |
| 3512qR | qPCR | qPCR primer of PL3512 | GCGAACTTCAGAAGCCACTT |
| 3518qF | qPCR | qPCR primer of PL3518 | CGACGCTCTTAAAGCCGATT |
| 3518qR | qPCR | qPCR primer of PL3518 | CGCGCACCTAGGTAAAGATG |

**Movie S1. Structural Alignment of Glycoside Hydrolase Domains of the Four Proteins.** The glycoside hydrolase domains of the four proteins were extracted based on Gene3D annotation results, loaded into PyMOL, and the aligned structures were displayed.

**Movie S2. Structural Alignment of CBM-like Domains of the Four Proteins.** The CBM-like (carbohydrate-binding module-like) domains were extracted from the four proteins based on the Gene3D annotation results. Except for PL3506, which contains one domain, the other three proteins each contain two of these domains. The aligned structures were loaded and displayed in PyMOL.

**Movie S3. Conserved Sites and Active Center of the Glycoside Hydrolase Domain of PL3506.** Within the structures, the active sites predicted by docking are marked in red, the sites that are both structurally and sequentially conserved within the glycoside hydrolase domains of PL3506 and a few other homologous proteins are marked in green, and the conserved sites mutated in the PL3506m mutant are marked in blue. These marked structures were loaded and displayed in PyMOL.

**Data S1.**

Whole-genome annotation information and proteome abundance results.

**Data S2.**

Genome accession numbers of 629 bacterial strains downloaded from NCBI.

**Data S3.**

Structural and sequence similarities among the glycoside hydrolase domains of the four proteins, some of their homologous proteins, and reconstructed ancestral proteins.

**Data S4.**

The Ka and Ks values between the four proteins and their respective homologous proteins.

**Data S5.**

The structural conservation index values between the four proteins and the homologous proteins.

**Data S6.**

Localization prediction information for the four proteins, their homologous proteins, and reconstructed ancestral proteins.

**Data S7.**

Gene–gene covariation, pairwise identity and evolutionary rate.

**Data S8.**

Partial geographical information of *Agarivorans* genus strain samples

**Data S9.**

Information on carbohydrate small molecules used for molecular docking.

**Partially expressed mutants and synthesized gene sequences**

>6g
GGTTGTTTAGATAAGTTTGGTCAAAATGCTTTAGTAGAAACCGCGGAAAAAGTGCAT
TCTGAAGAAGAGTTATTGGCGGTTACAGCCAAAGAGCTTAAAGCCTTAGAGCAAGG
TGCTATGCCCCAGCGTTCGCGCTTTAGTGGTTACACTGGAGGTCCGCAGCTAGAGGC
TACCGGCTTTTTCCGTACCGAAAAAATCGACGGTAAATGGTCGTTAGTTGACCCCGA
TGGCTACCCATACTTTGCTACCGGCCTCGACATTATTCGCCTAGCAAACTCATTTACC
ATTACCGGTATTGATTACGACCACTCAAAAGTGGCTGCGCGTAGCGATGACGATGTA
ACTCCTGAAGACTCTAAAGAAAAGCTCGAAATTAGCCAAGAAGCCTTTGATTCAGC
CTATGTTGCTAACCAAACTCGCCGAGACTTTTTTGATTGGTTGCCTAGCTACGACGA
CCCCTTAGCTGAGCATTACAGCTATATGCGTGAGCTGTTTGAAGGGCCAGTTGATCG
CGGCGAGATCTTTAGTTTCTACTCGGCAAATCTACAGCGCAAGTACGGTCAAGATGG
CGCCGATTACATGGGCAAGTGGCGCGAAGTTACCATGGACCGCATGCTTAACTGGG
GCTTTTCTTGTTTAGGTAATTGGACTGCTCCAGAATTTTACAGTAACGACAAAATTCC
TTATTTCGCCAATGGCTGGATTATTGGTGACTTTAAAACAGTGACTAGTGGCGATGA
CTTTTGGGCGCCACTGCCAGATCCTTTCGACCCGGTATTTAGAGAGCGCGCCGAAGC
AACGGTAAGCCAAGTTAAAGCCGAAATGCAAGGTTCACCATGGTGTGTGGGTATTTT
CATTGATAATGAGAAAAGCTGGGGCCGAATGGGCACCATTAATGGCCATTATGGTA
TTGCGATCCATACTCTAGGTCGCAGCGATGAAGAAAGCCCAACTAAGGCGGTATTC
ACTCAAGCCTTAAAAGACAAGTACGGCACTGTAGAACAGCTTAATCAAGCTTGGGG
GACGAACATCGACTCTTGGCAAGCAGTAGCTGGTGGCGTTAGTGATTTAGCCCATAA
CGAAGCACAGCTTAGTGACTACTCCATGTTATTAGAGTTGTACGCCAGCGAATACTT
CAAAGTGGTAAACGAGTCGCTTAAAGCTCAATTACCTAATCACTTGTATTTGGGGGC
TCGATTTGCTGATTGGGGCTGTAATCCAGAGGTGGTAAGAGCGGCTGCAAAACATGT
GGATGTTGTAAGTTACAACTACTACAAAGAAGGGCTACATCCAGAACCTTGGAAGT
TTTTGGCTGAAGTAGATATGCCGAGCATTATTGGTGAGTTTCATATTGGCGTTAAAG
AAGGTTTCTTCCACGCAGGTTTGGTGACGGCTAATGACCAAACCGAGCGCGGCGAG
ATGTTTGAAGATTACTTAAACTCAGTAATCGACAACCCATATTTTGTCGGCGCACAT
TGGTTCCAATACATTGATTCACCGATTACTGGGCGTTCGTTTGACGGTGAAAACTAC
AATGTGGGTTTTGTAGGCATAACCGATGTGCCATATCAGCCAATGGTTGATGCTGCT
AAGCGGTTAAATGGCGGCATGTATCAGCGCCGT

>6gM
GGTTGTTTAGATAAGTTTGGTCAAAATGCTTTAGTAGAAACCGCGGAAAAAGTGCAT
TCTGAAGAAGAGTTATTGGCGGTTACAGCCAAAGAGCTTAAAGCCTTAGAGCAAGG
TGCTATGCCCCAGCGTTCGCGCTTTAGTGGTTACACTGGAGGTCCGCAGCTAGAGGC

TACCGGCTTTTTCCGTACCGAAAAAATCGACGGTAAATGGTCGTTAGTTGACCCCGA
TGGCTACCCATACTTTGCTACCGGCCTCGACATTATTCGCCTAGCAAACTCATTTACC
ATTACCGGTATTGATTACGACCACTCAAAAGTGGCTGCGCGTAGCGATGACGATGTA
ACTCCTGAAGACTCTAAAGAAAAGCTCGAAATTAGCCAAGAAGCCTTTGATTCAGC
CTATGTTGCTAACCAAACTCGCCGAGACTTTTTTGATTGGTTGCCTAGCTACGACGA
CCCCTTAGCTGAGCATTACAGCTATATGCGTGAGCTGTTTGAAGGGCCAGTTGATCG
CGGCGAGATCTTTAGTTTCTACTCGGCAAATCTACAGCGCAAGTACGGTCAAGATGG
CGCCGATTACATGGGCAAGTGGCGCGAAGTTACCATGGACCGCATGCTTAACTGGG
GCTTTTCTTGTTTAGGTAATTGGACTGCTCCAGAATTTTACAGTAACGACAAAATTCC
TTATTTCGCCAATGGCTGGATTATTGGTGACTTTAAAACAGTGACTAGTGGCGATGA
CTTTTGGGCGCCACTGCCAGATCCTTTCGACCCGGTATTTAGAGAGCGCGCCGAAGC
AACGGTAAGCCAAGTTAAAGCCGAAATGCAAGGTTCACCATGGTGTGTGGGTATTTT
CATTGATAATGGGAAAAGCTGGGGCCGAATGGGCACCATTAATGGCCATTATGGTA
TTGCGATCCATACTCTAGGTCGCAGCGATGAAGAAAGCCCAACTAAGGCGGTATTC
ACTCAAGCCTTAAAAGACAAGTACGGCACTGTAGAACAGCTTAATCAAGCTTGGGG
GACGAACATCGACTCTTGGCAAGCAGTAGCTGGTGGCGTTAGTGATTTAGCCCATAA
CGAAGCACAGCTTAGTGACTACTCCATGTTATTAGAGTTGTACGCCAGCGAATACTT
CAAAGTGGTAAACGAGTCGCTTAAAGCTCAATTACCTAATCACTTGTATTTGGGGGC
TCGATTTGCTGATTGGGGCTGTAATCCAGAGGTGGTAAGAGCGGCTGCAAAACATGT
GGATGTTGTAAGTTACAACTACTACAAAGAAGGGCTACATCCAGAACCTTGGAAGT
TTTTGGCTGAAGTAGATATGCCGAGCATTATTGGTGAGTTTCATATTGGCGTTAAAG
AAGGTTTCTTCCACGCAGGTTTGGTGACGGCTAATGACCAAACCGAGCGCGGCGAG
ATGTTTGAAGATTACTTAAACTCAGTAATCGACAACCCATATTTTGTCGGCGCACAT
TGGTTCCAATACATTGATTCACCGATTACTGGGCGTTCGTTTGACGGTGAAAACTAC
AATGTGGGTTTTGTAGGCATAACCGATGTGCCATATCAGCCAATGGTTGATGCTGCT
AAGCGGTTAAATGGCGGCATGTATCAGCGCCGT
>PL3512M
ATGGTAGAAGTTATGAAATTTACAAAAAATAAAATTGCTGCGCTGTTATCGCTCACT
TTGTTGGGTGTTTACGGATGTGGCTCAACTCCTAGCTCGAGCGACGCCGAAGGCGCT
GTTGAAGATGTAGGCGGAACGATTCCAGATTTTGAATCTGCCGCTTTTTTCAAAAAA
GTTAAAAAAGACCATGCTAAAGCCGAAGTGGTTAGCGACCAAGGTGTTACTAGCGG
AAGTAGTGCGCTAAAAGTAAACTTTGATTCGGTATCCGAGGCTAACAAATTTAAGTA
CTGGCCAAATGTTAAAGTACACCCAGATTCTGGCTTTTGGAACTGGAATGCGAAAGG
CAGCTTAAGCTTAGACATTACCAACCCAACTGATAGCCCAGCAAACATCATTCTAAA
ATTGGCTGACAACGTGGGCGTAATGGGCTCTGGTGACAACCAATTAAACTACGCGG
TTAATGTACCCGCTGGCGAAACAGTGCCAGTTGAGATGTTATTTAACGGCACTAAAC
GTAAGTTAGATGGCTACTGGGGTGGCGAGAAAATCAACTTACGTAACATTGTAGAA
TTCCAAATCTTTGTGCAAGGTCCAATGGATGCACAAACTGTCATTATCGATAACTTT
AACTTAGTGGATGCTACCGGTGACTTTATTGAAGCCTCTGGCCAAGAAGTTAAAGTA
AGTGGTCCAATTCCAACAGTTGCTTCAATTACTAGCTTTGATGAAGGCCAACCTACC
TTTGTTGCTTTTGACCGCAGTGCAGCAGCAACGGTAACCGAGTTAAAAACCGACATG

GGCGGTTTGCTAGCTGTTAAATTAGCAGCGACTAACGCCTACCCTAATATCACTTTT
AAAGCGCCACAACCATGGGATTGGAGCGAGTACGGTGATTTTAGTTTAGCGTTTGAT
CTAGAAAGTAAAACAGACGAACCTTTACAGTTATTTGTTCGCGTGGATGACGCAGA
AAATGAAAACTGGGGCGGCACTGCAAACGGCGTAGTCGACAGTATGTCTAGCTACG
TAACGCTAGCTCCTGGTGACGACGGTACCTTCTACTTGCCACTAGGCCAAACCGGCA
GCCAAATCGTTTCTGGTATGCGTGCCGAGCCACCTAAAAAGTCTTACAACGCGCAAG
CGATTAGTTATGGCTGGGGTGAAAAGAGCTTAGATACGTCAAACATTGTATCTTTCC
AGCTATACCTGCAAAACCCAACTAAAGATGCAGAGTTCAATATTAAGAGTGTTCGA
CTTATTCCAAATATTGATGCAGATGCAACCCGTTATGAAGGTTTGATTGACCAATAT
GGTCAGTTTACTGGTTCTGAATGGCCTAAGAAAATTACCGAAGACGAAGAACTAGA
AACCATGGGTAAGCTGGCAAAAATGTCGCTTAAATCAACCAGCCAAATGCCAGGTC
GAAGCATCTATGGTGGCTGGGCCGATGGTCCTAAGCTGAAAGGCACAGGTTTCTTCC
GTACTGAAAAAGTAGACGGTAAGTGGTCGTTAGTAGACCCACAAGGCAACTTGTTC
TTCGCAACCGGTGTCGACAACATTCGTATGGACGACACAGTAACCATTACTGGCCAC
GACTACGACCACTCAAAAGTGGCTGCGCGTAGCGATGACGATGTAACTCCTGAAGA
CTCTAAAGAAAAGCTCGAAATTAGCCAAGAAGCCTTTGATTCAGCCTATGTTGCTAA
CCAAACTCGCCGTTCAATGTTTACTTGGTTACCAGAAGATGACGATGTATTAGCCGA
GAACTATGACTACGCAAATTGGGTGCATTCAGGGGCACTTAAGAAAGGCGAAGTAT
TTAGTTTCTATGGCGCTAACTTGCAACGTAAATATGGCGGTACTTTCTCAGAAGCTG
AAAAAGTGTGGAAAGACATCACCATAGATCGCATGGTTGATTGGGGCTTCACAACC
CTAGGTAACTGGGCTGACCCAATGTTCTACGACAACAAAAAAGTTGCTTACGTAGCA
AACGGTTGGATTTTTGGAGACCACGCACGCATTAGCACTGGTAACGACTACTGGGGT
CCTATTCACGATCCGTTTGACCCAGAGTTTGTAAACAGCGTTAAGGCGATGACTAAA
AAGCTAATGACAGAAGTAGACAAAAATGACCCATGGATGATGGGGGTGTTTGTCGA
CAACGAGATTAGCTGGGGTAACACCAAAAACGACGCAAACCACTACGGTTTGGTGG
TTAATGCGCTTAGCTATGACATGAAGAAGAGCCCGGCTAAAGCAGCCTTTACCGAG
CACTTAAAAGAGAAGTACTGGGCCATTGAAGATCTAAACACTAGCTGGGGCGTGAA
AGTTGCGTCATGGGCAGAGTTTGAGAAGTCATTTGATCACCGCTCTCGCTTAAGCAA
GAACATGAAGAAAGATTATGCCGAAATGCTAGAAATGCTTTCTGCTAAATACTTCTC
AACGGTTCGTGCTGAGCTTAAGAAAGTACTGCCTAATCACCTATATCTAGGCGCACG
CTTTGCAGACTGGGGTGTTACGCCAGAGATTGCCAAAGGTGCTGCACCTTACGTAGA
CGTGATGAGCTACAACTTGTACGCCGAAGACCTTAACTCAAAGGGTGATTGGAGCA
AGTTAGCAGAGCTAGATAAGCCAAGCATTATTGGTGAGTTCCACTTCGGTTCAACTG
ACTCAGGCTTATTCCACGGCGGTATCGTGTCGGCAGCTAGCCAGCAAGATCGCGCTA
AGAAATACACCAATTACATGAACAGTATTGCCGACAACCCTTACTTCGTAGGTGCTC
ACTGGTTCCAATACATTGACTCTCCAACAACGGGTCGTGCTTGGGATGGTGAAAACT
ACAACGTAGGTTTTGTTTCAATCACTGATACACCATACGTTCCACTGGTTGAAGCCG
CTAAGAAATTCAACCAAGACGTTTACATGCTTCGTTACAAAAAA
>PL3506M
ATGGAAACGCGCTTAAGGGCTGACTCAGCTAAAACTATATCTGAGCATGAGCAATC
AATTTTGATTTATGATTTTGCTGAGCAAATTCCTAAGGCTTTTAGTTTTAGCAACGTG

GACGCAGAACTGGTAAGTGAAGGCAACGGCATTACTACTGGTTCGCAAGCCTTAAA
AGTTACTACACATAGCAAAGAAAACTTTTACACCTCAATCTTTATCGAACCTGAGCA
GCCTTTTGATTGGAGTGCGCTTCCTAACTTTAGTTTTGCATTTGACGTCACAAACTTA
GGCCGTCGTTCAACACAGATTTTCATCAATATCTTCGACAAGCAGGGGCAAATGCAT
AGCCGCAGCATTAACGTGGCGGGCGGTTCTTGCAAAACCTATTTAAACGAGCTTAAG
GGTGAGTTTTTAAAGGGTGGGCTTAACTATGAATCGGGTTTTCGCTCTAACCCTGCA
GCCTGGGATACGCCATTTCACTATGCAACGTGGATGTGGGGCGAGATGAACATCGA
TTTAAGTGCTGTTGCCAAAATTGAACTAAGCATTCACGGTACATTGATTGATCATCA
GTTGGTGTTAGATAACTTTAGAGTGATTTTTACCCCGGAATGTAATCCCAATTTTCTT
AAAGGTTGTTTAGATAAGTTTGGTCAAAATGCTTTAGTAGAAACCGCGGAAAAAGT
GCATTCTGAAGAAGAGTTATTGGCGGTTACAGCCAAAGAGCTTAAAGCCTTAGAGC
AAGGTGCTATGCCCCAGCGTTCGCGCTTTAGTGGTTACACTGGAGGTCCGCAGCTAG
AGGCTACCGGCTTTTTCCGTACCGAAAAAATCGACGGTAAATGGTCGTTAGTTGACC
CCGATGGCTACCCATACTTTGCTACCGGCCTCGACATTATTCGCCTAGCAAACTCAT
TTACCATTACCGGTATTGATTACGACGTTGCTAACCAAACTCGCCGAGACTTTTTTGA
TTGGTTGCCTAGCTACGACGACCCCTTAGCTGAGCATTACAGCTATATGCGTGAGCT
GTTTGAAGGGCCAGTTGATCGCGGCGAGATCTTTAGTTTCTACTCGGCAAATCTACA
GCGCAAGTACGGTCAAGATGGCGCCGATTACATGGGCAAGTGGCGCGAAGTTACCA
TGGACCGCATGCTTAACTGGGGCTTTTCTTGTTTAGGTAATTGGACTGCTCCAGAATT
TTACAGTAACGACAAAATTCCTTATTTCGCCAATGGCTGGATTATTGGTGACTTTAA
AACAGTGACTAGTGGCGATGACTTTTGGGCGCCACTGCCAGATCCTTTCGACCCGGT
ATTTAGAGAGCGCGCCGAAGCAACGGTAAGCCAAGTTAAAGCCGAAATGCAAGGTT
CACCATGGTGTGTGGGTATTTTCATTGATAATGAGAAAAGCTGGGGCCGAATGGGC
ACCATTAATGGCCATTATGGTATTGCGATCCATACTCTAGGTCGCAGCGATGAAGAA
AGCCCAACTAAGGCGGTATTCACTCAAGCCTTAAAAGACAAGTACGGCACTGTAGA
ACAGCTTAATCAAGCTTGGGGGACGAACATCGACTCTTGGCAAGCAGTAGCTGGTG
GCGTTAGTGATTTAGCCCATAACGAAGCACAGCTTAGTGACTACTCCATGTTATTAG
AGTTGTACGCCAGCGAATACTTCAAAGTGGTAAACGAGTCGCTTAAAGCTCAATTAC
CTAATCACTTGTATTTGGGGGCTCGATTTGCTGATTGGGGCTGTAATCCAGAGGTGG
TAAGAGCGGCTGCAAAACATGTGGATGTTGTAAGTTACAACTACTACAAAGAAGGG
CTACATCCAGAACCTTGGAAGTTTTTGGCTGAAGTAGATATGCCGAGCATTATTGGT
GAGTTTCATATTGGCGTTAAAGAAGGTTTCTTCCACGCAGGTTTGGTGACGGCTAAT
GACCAAACCGAGCGCGGCGAGATGTTTGAAGATTACTTAAACTCAGTAATCGACAA
CCCATATTTTGTCGGCGCACATTGGTTCCAATACATTGATTCACCGATTACTGGGCGT
TCGTTTGACGGTGAAAACTACAATGTGGGTTTTGTAGGCATAACCGATGTGCCATAT
CAGCCAATGGTTGATGCTGCTAAGCGGTTAAATGGCGGCATGTATCAGCGCCGTTTT
AATAAGGCGAAAAAG
>PL3518M
GTGTTGTTTAAAAAGAGCAACTTAGCAATCCTAGTATCAGTTGTATTAGCTGGGGTA
TCTACATCAAACGTCATTGCAGATGACACTAAGCAGTCTAGTGAAAATGCAGCCACT
TCGGGTGATATGACCAGCGCGGCAACGCCGCTTGATTTCACCGAGCCAGCAGTATTA

GAAAAAATTACCAATAGTCATAGTCAGTTCTCGGTGCTTAAGAAAACCGCAGAGCA
AAGTAAAGATGGCTTAAAGATGAATTTTGATGCGATCTCTGAAGCCGAGGCGCAAT
CTAAATGGCCAAACGTAAAAATCCACTCTAAGGCAGGCCCTTGGGATTGGAATACC
AAAGGCGGCTTAAAGGTAGCCCTAGAAAATCCAGGCAGTGAAGACGTTCGCATCGA
AATGAAAGTGAGCGATAACATTGGCATTATGGGGTCGGCCGACAACCAAGTTGATT
TACCAATTATTTTGCCAGCGGGCAAAACTACCACCGTAGACTTTTTGTTTAACGGTA
CACAAATGAATATCGACGGCTACCGCGGTGGTGCTAAGTTAAACCTAAAAAGCATT
GCCGAAATCCAATTCTATTCAGTGGGGCCTATTGCTGCGCAAGAAGTGGTAATTCGT
GACATTAACTTTATTGAACGTACCGGTGATTTTGTAAAATCGGAAGCGCGTGAAGCG
GAAGTTATTGCTGCACCCATTCCAACTCTTTTAGCGTTGTCTGATTTTGATGATGGCT
CAAAAGGCATTGTTAGCAAAACTCATGGCACAACCATCACCTCAGTAAAGCGTGAT
GAGGGTAAAGGCCTTAAAATTGATTACTCCGCAGACGCGTCTTACCCCAGCGTAACT
TTTAGTGCCGATAAACCTTGGAACTGGTCAGAATACGGTGACTTTACCTTAGCCCTT
GATATTGAAAATATTGGTGATGCCGGCGCACAACTGTTTATTCGCGTAGACGATGAC
GTAAATGAGAAGCAAGGGGGTAGCGCAAATGGTGTTATCCATAGTCGTACTGGTTA
TGTTCAGTTGCCAGCAGGCGAAGCGGGTACTTACTACTTCACTTTAGAAGAGCTAGC
TAAAACGCTTGATTCTGGAATGCGAGGCGAGCCACCGAAGAAGTCTTATCAGGCGC
AAGCAATCAACTTTGGTTGGGGTGAGCAGAAGCTAGATCTAAGCAATATTGTTAGCT
TCCAACTCTATATGCAAGATCTGCAAAAAGACCTGAGTTTAGTTATCGACAACATTC
GCTTAGTGCCTAACTTGAGCGCCGATACCAGCCGCTATGAAGGTTTGCTGGATGAAT
TTGGTCAGTTTACCAATGAAGACTGGGCAGAAAAAATTCACTCAGCTGAAGAGCTG
CAGGCGCATGCAAAAGCCGATGTTAAGTTGATTGATTCGGCCAAACCTATGGACGA
CCGTACGCCATATGGTGGTTGGAAAAATGGACCTAAACTAGAAGCGACGGGTTACT
TTAGAACTGAGAAAGTAGATGGGAAATGGTCTTTAGTTGACCCAAGCGGTTACCTGT
ATTTTGCCACAGGTTTAGACAACATTCGCATGGATGATACCTACACCACCACCGGCG
TTGGCTTTACCGATTTGGTGGATGCTCGTTCAGTGGCTTCGCAGTTACGTAACAGCAT
GTTTACTTGGTTACCTAGCTATCAAGATGCCTTGGCGCAAAACTATCAATATTCCAC
CATGATTCACACCGGCCCGCTTGAGCATGGCGAGGTTTATAGCTTTTACAGTGCTAA
CTTGCAGCGTAAATATGCGCCTGACTCTCGCGATGAAGCGATTGCGGTTTGGCGCGA
CGTAACGCTGGCTCGTATGTTGGATTGGGGTTTTACCTCGTTAGGTAACTGGGCCGA
TCCAAGTTTTTACGGTAATCAAAAAGTTGCGTACGTAGCAAACGGTTGGATTGTGGG
CGATCACCAACGCATTAATACCGGTAACGATTACTGGGGGCCAATGCACGATCCTTA
CGACCCAGAATTTGTAGAGTCGGTGAAAACTATGGCTAAGCAAGTTGCCGCAGAAG
TAGAGCAAGACCCTTGGTGTATAGGCACCTTTGTGGATAACGAAATGAGCTGGGGT
AACACCGAGTTTGACGCTAACCACTATGCCTTAGCGATTGCCGCTTTACGTGCTGAT
GCTAAAGATAGCTTTGCTAAAGCTGCGTTTGTTGGCTTGTTAGAAGCGAAATATGCG
CAAGATATCCAGGCGCTGAACAAGGCATGGGGAAGTGAACTTAAATCATGGGATGA
ACTGGCTAAAGGCTATGTGCATCAAGGTGATTTGAACGACGCTCTTAAAGCCGATTA
CAGCATGTTCTTGGCTGACCACTCTGATCGTTACTTTGCGATTGTACAGCAACAGAT
GAAGCAGGTATTACCTAACCATCTTTACCTAGGTGCGCGTTTTACCGAGTGGGGGAT
CACTCCTGAAGCTGCGAATAGTGCTGCCCAATATGTAGATGTAATGAGCTACAACCT

CTACGGCAACGACATGTCGAAGGGCGATTGGAGCCACCTTGCTGAGTTGGATATGC
CAAGCATTATTGGTGAGTTCCACTTTGGTGCTACCGACAGCGGTATGTTCCACCCTG
GTCTAGTGGCTGCCGATACCCAACAAGGGCGTGCCGAGAAGTATGCCCACTATATG
GATAGTGTTATTGCTAATCCTTACTTTGTTGGTGCGCATTGGTTCCAGTACCTAGATT
CGCCAACTACTGGTCGAGCTTGGGACGGAGAAAACTACAACAATGGTTTTGTCACTG
TAGCTGACTCGCCCTATGAAAAACTGGTAGCAGGAGCAGCAGAAGTTAACCGTAAG
CTGTATCCACAGCGTTACCCAGAGCTGGTGAAG
>PL3511M
GTGCAATCTGCTAGGTACAAGGAATGGGCAGGAAATATGAAGATTAAATTTTTATCT
GCAGCAATTGCTGCAAGCTTAGCATTGCCATTAAGTGCTGCTACTTTAGTCACCTCTT
TTGAGGAAGCCGACTACAGCAGCTCTGAAAACAATGCTGAATTTTTGGAAGTGTCTG
GAGATGCCACTTCTGAAGTTTCAACAGAACAAGCTACTGATGGTAATCAATCGATTA
AAGCGTCTTTTGACGCGGCTTTCAAACCAATGGTTGTTTGGAACTGGGGAAGTTGGA
ACTGGGGCGCTGAAGATGTTATGTCAGTAGATGTTGTTAACCCTAACGACACTGACG
TTACCTTTGCTATTAAGCTAATCGATAGTGATATTCTTCCTGATTGGGTAGATGAGTC
TCAAACCTCATTGGACTACTTTACGGTTTCAGCTAATACCACGCAGACCTTTAGCTTT
AACTTAAATGGCGGGAACGAGTTCCAAACTCATGGCGAAAACTTTAGTAAAGATAA
AGTTATCGGTGTGCAGTTCATGCTCTCTGAAAACGATCCTCAAGTGTTGTACTTTGAC
AACATTATGGTTGATGGCGAAACAGTCACTCCGCCACCAAGTGATGGTGCAGTGAA
TACACAAACCGCGCCTGTAGCCACCTTAGCGCAAATCGAAGACTTTGAAACCATTCC
AGATTACTTACGACCTGATGGTGGGGTAAACGTTTCAACTACTACTGAGATTGTGAC
TAAAGGCGCTGCAGCAATGGCTGCTGAGTTTACTGCAGGTTGGAACGGTTTAGTGTT
TGCAGGTACTTGGAATTGGGCAGAACTAGGTGAACACACCGCAGTCGCCGTTGACG
TTTCAAATACTAGCGATAGCAATATCTGGTTGTACTCACGTATCGAAGACGTAAATA
GCCAGGGCGAAACTGCGACTCGCGGCGTATTGGTTAAAGCTGGCGAATCGAAAACC
ATTTACACCAGCTTAAATGATAATCCTTCATTGCTTACTCAAGATGAGCGTGTGTCA
GCTTTAGGTTTACGTGATATTCCAGCTGACCCAATGAGCGCTCAAAATGGCTGGGGT
GATTTTGTTGCTTTAGACAAATCTCAAATTACCGCTATTCGTTACTTCATTGGCGAAT
TAGCCAGTGGTGAGACTAGCCAAACACTTGTGTTTGATAACATGCGCGTGATTAAAG
ACCTTAACCACGAATCAGCCTATGCAGAAATGGCCGATGCTATGGGGCAAAACAAT
TTAGTCACTTATGCAGGTAAAGTTGCCAGCAAAGAAGAGTTAGCTAAGTTAAGTGAT
CCAGAAATGGCTGCTTTGGGTGAGTTAACCAATCGCAATATGTACGGTGGTAACCCA
GATTCGTCGCCAGCTACAGACTGTGTGCTAGCTACGCCTGCCTCGTTTAACGCTTGT
AAAGACGCTGATGGTAATTGGCAATTGGTCGACCCTGCTGGTAATGCGTTCTTCTCA
ACCGGTGTTGATAACATTCGTTTGCAAGATACTTACACCATGACCGGCGTGTCGAGT
GACGCCGAATCTGAGTCTGCACTTCGCCAGTCAATGTTTACAGAAATTCCAAGTGAT
TATGTAAATGAAAACTATGGCCCTGTGCATAGTGGACCTGTTTCTCAAGGCCAAGCT
GTAAGTTTTTACGCTAATAACTTAATTACCCGCCACGCTAGCGAAGACGTATGGCGA
GACATTACTGTTAAGCGCATGAAAGACTGGGGCTTTAACACCTTAGGTAACTGGACC
GATCCTGCGTTGTATGCAAACGGTGACGTTCCTTACGTGGCAAATGGTTGGATTTTT
GGAGACCACGCACGCATTAGCACTGGTAACGACTACTGGGGACCACTTCCTGATCC

GTGGGATGCTAACTTTGCTACCAATGCCGCCACAATGGCTGCAGAGATCAAAGCTCA
GGTTGAAGGCAACGAAGAGTACTTAGTGGGTATTTTTGTTGATAACGAAATGAGCTG
GGGCAATGTCACTGATGTTGAAGGCTCTCGTTATGCGCAAACGCTAGCGGTGTTCAA
TACCGACGGCACTGATGCAACAACTAGCCCTGCCAAAAATAGCTTTATTTGGTTCCT
AGAAAACCAGCGTTATACCGGTGGAATTGCTGACCTAAACGCAGCCTGGGGAACCG
ATTATGCGTCTTGGGATGCGATGCGCCCAGCGCAAGAGTTAGCTTATGTGGCTGGCA
TGGAAGCTGATATGCAGTTCCTTGCTTGGCAATTTGCTTTCCAATACTTCAACACCGT
AAACACGGCATTAAAAGCTGAGTTACCAAACCACTTGTACTTGGGCTCTCGCTTCGC
AGATTGGGGACGTACTCCTGATGTAGTAAGTGCTGCTGCGGCTGTTGTTGATGTAAT
GAGTTACAACATCTACAAAGACAGCATCGCAGCTGCCGATTGGGATGCTGATGCTTT
AAGTCAAATCGAAGCCATTGATAAGCCAGTAATTATTGGTGAGTTCCACTTCGGTGC
GCTTGATAGCGGTTCGTTTGCAGAAGGTGTAGTAAATGCCACTTCGCAACAAGATCG
TGCAGACAAAATGGTTAGCTTTTACGAATCAGTAAATGCCCATAAAAACTTTGTAGG
TGCGCATTGGTTCCAATACATCGATTCACCATTAACGGGTCGTGCATGGGATGGCGA
AAACTACAACGTTGGTTTTGTTAGCAATACTGACACGCCATATACATTGATGACAGA
TGCTGCGCGTGAGTTTAACTGTGGTATGTACGGCACTGACTGCTCTAGCTTAAGCAA
TGCTACTGAAGCTGCTTCGAGAGCGGGTGAGTTGTACACCGGTACCAATATTGGTGT
TAGCCACTCTGGCCCAGAAGCGCCAGATCCAGGTGAGCCAGTAGATCCACCAATTG
ATCCGCCAACACCACCAACGGGTGGCGTAACTGGCGGTGGCGGTAGCGCAGGTTGG
TTATCGCTACTAGGTTTGGCCGGCGTATTTTTACTAAGACGTCGTAAAGTG
>Vsp6
ATGACACCAACCATTAATGATGTGGTGAAACACTCAAAGCATGATAGCTCCGTGGCT
CTGTTTGATTTCTCCACCGAACAACTACCCAAGGCTTTTCGCTTTAATAATATTGATG
CATCAATGACGTCACAGGCGAAGCTTAAAATCCAGTGTCATAGTGCAGAGAACATG
TATACGTCAGTGTTTCTTGAGCCTGAGCCTGGAGAAAAATGGGACTGGAGCCAAAT
GCCAGAGTTTTGCTTTGCTTTTGACGCTCACAATCTACGCTCCAGGTCCACTCAGGTT
TTTATCAATATTTTTGACTCTAAAGGACAAATGCATAGTCGCTGCGTCAATGTCACT
GGTGACACTGATACTAGCTACCTAGTAGAGCTTAAAGGTGAGTATTTAAAAGGCAA
CACCAACTACTATTCAGGTTTTCGTTCCAACCCTGCTCCTTGGGATAGTCCATTTGTT
TATGCCACTTGGATGTGGGGTCTAATGAACATCGACCTATCAGATATTGTGCAAGTC
GAACTTAGTATTCACGGCACACTCATTGACCATGAGCTTGAACTGTCCAATTTCCGT
CTCATGCTCAGCCCAGAGATAAACCCTAGCTACCTTTCCAACATTATTGACTGCTAC
GGTCAAAACGCAGGTTTTGAATACCCAGAGAAAGTCCACAACGATCAAGAGCTGAC
AGAGTTTACTGAGCGTGAACTTCAAATGCTTAAAGAAGGTGCGATGCCTGACCGCTC
TCGCTTTGGCGGATATAAAGAAGGTAAACGTTACGAAGCGACTGGTTTTTATCGCAC
TGAGAAAATCGATGGTAAATGGTCTTTGGTTGATCCAGAGGGTTACCCATATTTCGC
GACAGGAATCGATATCATACGCCTTGCGAACTCCTACACCCAAACTGGCGTAGATTA
CGACCATAGTAAAGTTGAACAGCGCTCCCCTGATGACTTAACACCTGAGGACTCGAT
TGAAAAATTCGAAGTTTCAATGGAAGCCAAGCAAACCGCATTTGTCGGTAGCGATG
TTCGTCGTAACTGCTTCCAATGGTTGCCAGACTACGACGATGAGTTGGGAGAGCACT
ACGCCTACATGCGTGAGAATTTTGAAGGTGCCCTTGATCAAGGCGAAACTTTCAGTT

TTTACGCCGCCAACTTACAACGTAAATATGGCAAGCAATACATGGATAAGTGGCGA
GAAGTCACCATGGATCGCATGCTTAACTGGGGCTTCACCTCGTTAGGTAACTGGACT
GCTCCTGAATTTTACTCCAATGAAAAGGTACCATTCTTCGCAAATGGGTGGATCATT
GGTGAATTTAAAACAGTAAGTAGCGGTGACGACTTCTGGTCTCCTCTTCCGGACCCG
TTCGATCCACTATTTAAAGAACGTGCAGAAGCTACCGTTAAGCAAGTTCGCGAAGA
GATCAAAGATACGCCTTGGTGTGTGGGTATCTTTATCGACAATGAAAAGAGTTGGGG
TCGTATGGGCACCATCGAAGGTCAGCACGGTATTGCAATCCATACCCTAAGCCGTGA
CGCGAATGAATGTCCTACAAAAGCAGAGTTTATGAACGTTCTGCGTGATAAGTATGG
TGATATTGAATCACTCAACGCTCGTTGGGGTACGGAAATCGCTTCATGGGAAGCGCT
AAGCCACGGTGTCAAAGGTTTAGCTAACAACGAAGCGCAGCTTGAAGACTATGGCA
TCCTATTGGAAGCCTACGCCTCTCAGTACTTCAAGATCGTTCGTGAAGCGCTAAAAG
TAGAGCTGCCAAATCACCTCTATTTGGGCTGCCGCTTCGCTGACTGGGGTATGACTC
CAGACGTAGTGCGCGCTGCAGCAAAATACTGCGATGTCATTAGCTACAACTACTACA
AAGAAGGTCTACATCCTCAGCCATGGAGCTTCCTATCTGAAGTAGATATGCCAAGCA
TCATCGGTGAGTTCCATATTGGCTCAAAAGATACGGGGTTATACCACCCTGGCCTAG
TGACCGCCGGTAATCAACAAGAGCGTGGTGAGATGTATGAAGCGTATATGCACTCA
GTTATCGACAACCCATACTTTGTTGGCGCACACTGGTTCCAATACATCGACTCACCC
ATTACTGGTCGCTCATACGATGGCGAAAACTACAACGTTGGGTTTGTCTCAATTGCA
GATACTCCATATGAACCTATGGTAGAAGCCGCGAAACGTCTACATAGCTCTATGTAT
AAACGTCGTTATAAATAG
>Vsp12
ATGAAAAAGACGAAAATTGCACTTATTCTATCAAGCTTGCTCATTGGCGCTTACGGT
TGCCAGTCGACGACGGGGTCTCAGCCTGAAGCTGCGACGGAAAGTGAGGAACAAAC
AAGTGTTGCTATCCCAGATTTTGAATCTGATAGCTTTTTTAATAATGCACAGGCATCA
CATGCAAAAGCGGAGAAGGTTTCGGGTGTTGGAGTAACTTCAGGTGACTCGGCACT
AAAAGTGACATTTGATTCCGTCTCTGAAGCCAACAAATTCAAATACTGGCCGAATGT
GAAGTTCTTCCCCGAAGGCGGAATGTGGAACTGGAACAACAAGGGCAGTCTAAAGG
TTGACGTCACGAACCCAACTGACAATCCAGCCACCTTCATCTTTAAAATCGTAGATA
ACGTTGGCCAAATGGGCGCGGCGACGAATCAGTTGAATTACGCGGTAACAATCCCT
GCCAATAGCACTGAAAAAGTCGAGATGCTATTTAACGGCGGAAAACGTGGTCTAGA
CGGTTATTGGGATGGTCAGAACATCAATCTTCGTAAACTTGCAGAGTTACAAGTCTT
TGTCCAAGGGCCGATGGACAAACAAACCATCATCTTAGATAACCTTGAATTGGTTGA
TGCGACCGGAGACTTCATTTCTGCTGAAGCGCAGGTCGTTGAAGCGGGTCCGATTCC
GACCTTGAAGTCGATTACCAGCTTCGCGTCTGGAGAAAAGCACTATGTATCCGATCG
CAGTGTGTCTACTTCTATTGAGAAAGTGAAGTCAGACGGTGGTCTGGGTCTACAAGT
GAAGTTTAAAGCCGATAACGCGTATCCCAACGTCACGTTTATGCCAAATCGCCCTTG
GGATTGGTCAAATGAAGGCAACTTTAATCTTGCTTTTGATATAGAAAACCCAACCAA
TGACCCAATTCAAATGTTTGTGCGAGTAGACCAGGCTGAGAATAAGAACTGGGGAG
GCACGGCAGATGGCGTACAAGACAGTATGTCGAACTACGTAACCTTGTTGCCAAAC
GAGAAAAGCACTTACTACATGACACTGAGTCAGCTTGAAGGGCATATCGTTTCCGG
AATGCGCAGTGAGCCACCAAAGAAAGCATATAGCGCACAAGCGATCAGCTACGGTT

GGGGTGAAACAAGCTTAGATTTATCGAACATTGTCTCAATCCAGCTCTACTTACAAA
ACCCTACCAAAGACGCAACCATTGTGGTTGATTCAATTCGCCTGGTTCCTAACCTTG
ATGCTGATGCCACACGCTATGAAGGTTTGGTCGATCAGTATGGTCAGTTTACAGGCA
GTGATTGGACAGAAAAGATCCATAACGATGAAGAGTTGCGAGAATCAGGTAAAGCA
GAACTTGCAGAGCTAGGCAATGCTAAACCGATGTCCGATCGCAGTAAGTTTGGTGGT
TGGAATGCGGGCCCGCGTCTTGAAGCGACGGGTTTCTTCCGCGCAGAAAAGCTTGA
GGGCAAATGGACTTTGGTTGATCCAGAGGGCTACCCGTTCTTTATGACGGGACTGGC
GAATATTCGTATGGATGACACGGTTACGATTACCGGTGCCGATTACGCTAACCCAAA
AACGAAAGAGGGTCGCTCAATTGCTTCTCCGCTGCGTGAGTCGATGTTTACTTGGCT
ACCAGACTATGATGACGAGTTAGCAATCAACTACGACTATGCTGGATATGTTCACTC
GGGTGCGCTTAAGAAAGGGGAAGTATTTAGCTTCTACCGTGCAAACTTAATGCGTAA
ATACGATACCAATCAGCAAAAGGCGTTAGAAATCTGGCGAGATGTGACGCTTGATC
GTATGCAAGATTGGGGTTTCACCACGTTGGGCAACTGGATTGATCCGATGTTCTTCG
GTACAGAACGTGTGGCTTATGAAGCCCATGGTTGGATTGCTGGTGATCACAAACGA
ATCAGCACAGGTAATGATTACTGGGGGCCAATTCATGACCCGTTCGACCCAGAGTTC
AAGCAGAGCACTCGTGCTATGGCAGAAGGGCTTGCCAAACAGGTAGATAAGAACGA
TCCATGGCTACTCGGTATCTTTGTTGATAATGAAATCAGCTGGGGTAACGTCATGAA
CGAAGCCAACCACTTTGGTTTGATCGTGAATGCCTTGAGCTATGACGCAAAAGAGA
GTCCAGCTAAAGCCGCTTTCTCTCAACATTTGAAAGAGAAATACACCACAATCAATG
CTCTCAACAAAGCGTGGGGTACGACTCAAGTCAGCTCGTGGGAGGAGTTTGACAAG
TCATTTGATTATCGAAGCAGTTTGAATCCAGGCATGAAGCGTGACTACTCAGACATG
CTAACGTTACTGGCCGATAAATACTTCGCGACAGTCGACGCAGAGCTAGAACGTGT
ATTACCAAACCATATGTATCTTGGAGCCCGTTTTGCTGACTGGGGGGTTACGCCAGA
AATCGCGAAAGGCGCTGCAAAATATGTAGACGTCATGAGTTACAACCTATATGCCA
ACGACCTGAACGATTCGAAAAAAGCACACTTTAGAAAGTGGTTAGCGGAGTTAGAC
AAACCAAGCATCATTGGCGAGTTCCACTTCGGTTCTACTGATACGGGTCTGTTCCAT
GGTGGTATTGTCAATGTCGCTAATCAAAGTGAGCGTGCGAAGATGTACACCCACTAC
ATGCAGAGCGTCGTTGATAATCCGTATTTTGTCGGCGCACACTGGTTCCAATATCTT
GACTCTCCAGCAACAGGGCGAGCTTGGGATGGCGAGAACTACAATATCGGTTTTGTC
ACGATCGCTGACGAGCCATACGTTGAGTTGATTGAGGCGGCAAAACAATTCAACGC
AAACCTATATAACAATCGTTTCAAATAA
>Vsp11
ATGAACGTGAATAAAACTCTGACAGCTCTCCTGATTGCAGGAATGGCTGCTCCTACG
CTTGCTTCAACCCTTATCACTTCGTTTGAAGATAGCGAAACCCATGGTTATGAGTAT
GGACACTTTGGTAGTTCAACACATGTTATTACGTCTGATGTTCAAACTGACGGCGAA
AAATCGATGCAAGCCAATATGGGTGCCAGTTATCAAGGTGGCGTAAAAATATACGC
ACCGAATAGTTGGAATTGGACGGATAAGAAGGAGCTCGTTTTTGATATTGTTAATAA
TACCAACGAAGAGATCAATTATGGCGTGAAGATCCTTACCAACTACGTTTGGGATGA
TGCTAATGCTCTGTCCGAGTACTTTAATGTTCCAGCAAACACGGTACTTGAAAACGT
TGTGATGTCGTTAGACTCGACGGAGTATGGTTCAATTGGTGCTTCTTATAACAAAGC
GATGGCTATGGAAATCAACTTCTTTGCCGCTGAGTCACCAAACACAACCGTTTACAT

TGATAATATTCGTGTCTCTGATGGAGCGACGGAGCCAACCCCACCGCCAACAGATCC
TGAGGAAGCCAGTGGTCGTGTTTCATCAGCACCGGTTAAGACACTGCTGCAGCTTGA
AACGTTTGATGAAATTACCGACGCCATCGAAACTTCAGGCTCTGACATAGCGTTAAT
TGATGAAGGGGTGAGTATTGGTGACACAGCACTGAAAGCCACTTTTAACAATAGTT
GGAGCTCAGTGAAACTGCATGGGAACTGGGATCTATCCGCGCTTGGTGACCATCTAG
CAATTGCGGTTGATGTAACCAATCCGACCAATACAGGACTGTTCTTATACTCACGAG
TAGAGGATAACAGCGGTAATGATGCGTCGGGAATTGCTCGTGGTAAGTACATCCCA
GCCAACAGCACTCAGACTGTTTATGTGAGCGTTAAAGACACGCCTGCAATCAGTGAT
ATTGTCAATACGCTGGGCATGCGCGAACTACCAGCGAAAGCTTTGAGTGAGGGGTG
GGGTGATTGGCAAGAATTAAACCTAGCCGCAATTGAGGCTGTGACATTCTTTGTTCC
TGACTTGGCTGCAGGCACGACAGAGCAGCAGTTTATCTTTGATAATGCCCGTGTGAT
CAAAGATCTAAATCATGTATCTGCTTACGAAGAACTAGTGGATGTGCTTGGCCAAAA
TAATCAATACGACTTCTACGCTAAGTTAGAGGAAAGTGAGGATTTGGAGGGGCTGG
GTGAACGTGAAACACAACTGCTAGGCAAACTACTTAATCGAAACCAATACGGCGGA
GCGCCCTCTGGTAGTAATATTATTGCCGATCAAGACTGTAAACTGTCATCGCCCGCG
TCATTTAATGCGTGTAAAACCGCAGATGGCAAGTGGTATCTCGTAGATCCTGATGGA
CATGCTTTCATTTCGACAGGTTTGGCAAACATACGAATGACAGACACCTACACGTTT
ACCGGAGAGTCGAGCACGACCCCTTCGGACGTCCGTAAGTCGATGTTTACTGAGATT
CCTGCGAATCACCGCAAAGAGATGGACCGGTCCACAGTGGCCCTGTCGAAAAAGG
TGAAGGCGTAAGCTTTTATGCTAACAACATCGATGCTCGCCACGGTGGTGAGGAAG
CATGGCAAGATATTACAATCAAGCGAATGCAGGATTGGGGCTTTACTTCTTTAGGTA
ACTGGACCGATTCTGCGTTTTATTCGAAAGCGGCCGCAGCGAATATGCCTTACGTGG
CAAATGGCTGGGTACTTCACCATGAGACATCAGAAAACCCAATTAACCGTATCGGTT
CCGGGTATTGGGGGCCGATAGCTGACCCATTTGATCCTAACTTTGCGCTAGCTGCTA
AGAAGATGGCAGAGCAAATCAAGCTAGAAGTTACTGGACATGAAGCCACATTGATG
GGAATTTTTGTTGATAACGAAATTAGCTGGGGCAACTGTCTCAATGGGGATGATGCA
TCCTGTTATGAGCAGACGCTAGCCGCAATGAACACCGATGCGAGTACTAGCCCAGC
CAAAAATTCGTTTATGTGGTTCTTAGAAAATGGTTACGGTCATTCAAAATCTATCGA
GCAATTTAATGCAGCGTGGGGGACATCCTTCGCTTCATGGACTGACTTGAGAGGTGC
TCAGAACTTTAAATACTCAGCGGGTATGCTGGGAGATTTGAAGTCTCTAAACTGGCA
GTTCGCTAATAAGTACTTTGAAGTGGTTAACCAAGCGGTTAAAGCTGAGTTTCCAAA
TAACCTTTATTTAGGCGCGCGTTTTGCGGATTGGGGTCGCACACCAGAAACGGTTGA
TGCTGCGAAAGGTCATGCAGATGTCGTGAGTTTCAACATTTACAAGGACAGCATCAC
GTCGCAACATTGGCAGTCAGATGTCTTAGAGCAAATTGAATCTCTAGATTTCCCAGC
GATCATTGGTGAGTTCCATTTTGGTGCTCTAGACAGTGGTAGCTTTGCAACGGGAAT
CGTCTCGGCTGATTCACAGCAAGATCGCGGAGATAAGTACACCGCGTACATGGAGT
CGGTGCTAGATAACGATAACTTTGTTGGCGCTCATTGGTTCCAATACCTCGACTCAC
CAGTGACAGGTAGAGCTTGGGACGGAGAGAACTACAATGTCGGATTTGTTAACGTG
ACAGATACACCATACAGCCACCTTACTGACGCAGCACGTATTGTTAACTGTGAACTT
TATGGCGATGATTGTTCAGCATTGAAAGGAGATAGTGAAGAGCGCTCTGCCAGAGA
TTCAGGGGCACTTTACGATGGTAAAAATATCGGTATCACTTCAGGTCAGTTAAATGG

CATGGAGACCATTGATGGAATAGACCCAGAAGAACCGGTGGATCCGCCGACTGATC
CAGATGAGCCTCAGTTACGCACTGGTGGTTCTGCTGGTGGGCTATTTATCACTCTAA
TGGCAATAGCGGGCTGGTTGCGTAGAAAGTACGTATCCTAG
>Vsp18
GTGCGTTTCAACAAAAATACTATCGCGCTCGCTATCATTGCCAGTACTCTAGCTACG
GGTAGCTTAGCAGCAGCCAAAACACCAGAAGCATCGACAAATGAAGCGACTCAACA
AGATATGAGTAGCGCGGTGCAACCCGTTGTCATGACAGATGAAGGGTTTCTATCGA
AAACAGCAAATACACATACTTCATACCAAGTTGCTGGTGAGAATCTAGAGATTGTAT
TTGATGCAATATCAGAGTCCGAGGCCAACTCTAAATGGCCCAATATGAAGTTCCGTC
CAGAGTCGGGCTCTTATGACTGGAATACCAAAGGTGGGTTGCAACTGACACTTGAA
AACCCAGGCGATAAAGAAGTTCGTATCGAAATGAAGGTGGCTGATAATTTAGGCAT
CATGGGGGCTGCAACGCATCAACTGGATTTACCCATTTATCTCGCAGCGGGTAAAAC
CACGACGGTCGATTTCCTGTTTAATGGCGCGGAAATGAACATTGAGGGCTATCGTGG
CGGTAGTGAATTGGATCTGCGTAATATCGCAGAGTTTCAGTTCTACTCTGTAGGACC
TATCGCTGAGCAAAAAGTGGTGGTCCATGGTATCGATTTTATTGAACGCACGGGTGA
CTTTGTTGTGTCGGAAGCTCGTCAAGGGCAGGTGATCGAGGCTCAGATTCCGACACT
GCTTACAGTCACTGATTTTGAACAAGGTATTGATCAGATCGTTGAACGACATACGGG
CTCGAATGTTGATATCGTCGAAACTTCAACAGGCTCTGGTATTAAAGTTCACTACAC
TACCGATGATGATTATCCAACGATCAAGTTCTCTGCCGGTACAAATGGTGAGGCATG
GGATTGGTCTGAATTTGGCGATGTTGCACTTGCGTTTGATGCGAAGAACCTGGGTGA
CTCGGGTATGCAGTTGTTCGTTCGTGTCGATGATGCTTTGGATGAAAAGTTGGGCGG
TACGGCAACAGGGGCAGTGAACAGTCGCACCGGTTACGTGCAAGTCCCAGCCAATT
CCGAAGACAAATACTATTTCACTTTTAAAGACCTCGCTGAAGGTCTTGATTCTGGAA
TGCGTGGCGAGCCACCTAAAAAATCATTCTCTGCAAGCCAAGTTGTTTTTGGCTGGG
GTGAAGCTGAGCTTGACCTATCAAATATCGTCAGTGTTCAGCTTTACATGATGAATC
CTCAAGAAGAAGCGACTTTGGTGATTGATAATCTATCACTGATTCCAAACCTGAGCA
CTGATACAACGCGCTATGCAAACCTTCTTGACGAGTTTGGTCAATACCAAGAAGAGT
CATGGCCTGAGAAAGTCACTGATGTGGCACAGCTAAAAGAGCAGGGCAAAACTGAC
AAAAAGCTGCTAAAAAAAGCGGCCTTAATGGATGATCGCTCAAAGTTCGGCGGTTG
GGCTAAAGGACCAAAATTAGAAGGTACTGGTTACTTCCGCACTGAAAAAGTAGATG
GTACATGGGCACTGGTTGACCCAGAGGGCTACCTATATTTCGCTACGGGTGTAGACA
ATATCCGCATGGATGATACCTATACGGTAACGGGGATGGACTTTGCTGACGCTGTCG
AAGCAGATACTAAAGGTATGCGTCCAAGTCAGGTCGCAACCGCACGTTATGTGGAT
GACAAACGTGAGCGTGTTGAAGCTTCAACACTGCGTCGTGGTATGTTTGATTGGTTA
CCTGATTTCAACGATCCGTTGGCAGATAACTACAGTTACACTCAAATGGTACACACA
GGACCACTTAAGCACGGCGAACTCTTTAGTTTCTACTCTGCTAACTTGCAGCGCAAA
TACGACACAGATACCGCCCAGGAGGCGATTGATATCTGGAAAGATGTCACTCTGGC
TCGTATGCAAGATTGGGGCTTTACGTCATTAGGTAACTGGACCGACCCAATGTATCG
TAAGAATGGTAAGGTGCCTTATACGGCTCACGGTTGGATTACGGGTAATCACCAAA
GAGTAAGCACTGGTAATGACTACTGGTGGGCGATGCACGACCCGTTTGATCCTCAAT
TCCGAGTGTCAGTGGCGACTATGGCCAAAGCATTGGGTGAAGAGGTGGATAACGAC

CCTTGGTGTATTGGTTACTTCGTTGATAACGAACTAAGTTGGGGTAATACCGTCAAC
GACACGAACCATTACGCTTTAGCGGTTTCTGGTTTACGTGAGAGTGCAGAATCAAGC
TCAACAAAAGCCGCTTTTGATGCGTTGCTAAAAGAAAAGTACGGCTCAGTTGAGAA
GTTCAACCAAGCTTGGGGCACAGACGTCGCGTCTTGGAATGAATTTGCCAAAGGATT
CAACTATCAAGGTGAATACACGGACACGGTTAAAGCTGACCTGTCGATTCTATTAGA
TGCATTTGCTGATGAGTTCTTCGCTGTCGTCAGTGAAGAGATGGAGAAAGTGTTACC
TAATCACCTCTACATGGGCGTTCGTTTCTCTGACTGGGGCATCACACCGGAAGCGGC
GACAGCAGCCGCGCGCTATGTTGATGTGATGAGCTACAACTTGTATGCCACTGACTT
GAATGCAAAAGGGGATTGGAGTCGTCTTCCTGAGCTAGACAAACCAAGTATTATTG
GTGAGTTCCACTTTGGTTCAACCGACTCAGGTTTGTTCCACCCTGGTATTATCAGTAG
TGACGACCAGAAAGGTCGCGCAGAGTCTTACGCTAAGTACATGGAAAGTGTTATCG
ACAACCCATACTTCGTAGGTGCACACTGGTTCCAATACATGGACTCACCGGTAACAG
GTCGTGCGTGGGATGGCGAAAACTACAACGTGGGCTTTGTAACGGTAACGGACACA
CCATACGAGCCTTTGGTAGAGTCGGCTAAGGAGATTAACCGCAATCTTTATAACCGT
CGCTTCGGTTCGTTGAACTAG
